# Ecophysiological differentiation between life stages in filmy ferns (Hymenophyllaceae)

**DOI:** 10.1101/2021.03.12.435213

**Authors:** Joel H. Nitta, James E. Watkins, N. Michele Holbrook, Tristan W. Wang, Charles C. Davis

## Abstract

Desiccation tolerance was a key trait that allowed plants to colonize land. However, little is known about the transition from desiccation tolerant non-vascular plants to desiccation sensitive vascular ones. Filmy ferns (Hymenophyllaceae) represent a useful system to investigate how water-stress strategies differ between non-vascular and vascular stages within a single organism because they have vascularized sporophytes and nonvascular gametophytes that are each capable of varying degrees of desiccation tolerance. To explore this, we surveyed sporophytes and gametophytes of 19 species (22 taxa including varieties) of filmy ferns on Moorea (French Polynesia) and used chlorophyll fluorescence to measure desiccation tolerance and light responses. We conducted phylogenetically informed analyses to identify differences in physiology between life stages and growth habits. Gametophytes had similar or less desiccation tolerance (ability to recover from 2 d desiccation at -86 MPa) and lower photosynthetic optima (maximum electron transport rate of photosystem II and light level at 95% of that rate) than sporophytes. Epiphytes were more tolerant of desiccation than terrestrial species in both life stages. Despite their lack of greater physiological tolerances, gametophytes of several species occurred over a wider elevational range than conspecific sporophytes. Our results demonstrate that filmy fern gametophytes and sporophytes differ in their physiology and niche requirements, and point to the importance of microhabitat in shaping the evolution of water-use strategies in vascular plants.

## Introduction

The movement of plants from water to land was one of the most important evolutionary events in the history of the earth, fundamentally altering the global carbon and water cycle and setting the stage for modern terrestrial ecosystems (Graham 1993; Kenrick and Crane 1997). The transition from aquatic to terrestrial growth was complex, requiring a suite of evolutionary adaptations to survive in deadly dry habitats bombarded with radiation (Waters 2003). Lacking mechanisms to regulate water loss, the earliest land plants were likely poikilohydric, i.e., they existed in a state of equilibrium with the humidity of the external environment (Raven 1999). Therefore, one particularly important early adaptation that enabled plants to survive on land was desiccation tolerance (DT), the ability to lose all metabolically active water, reach equilibrium with atmospheric humidity, and recover upon rewetting (Mishler and Churchill 1985).

Compared with the ubiquitous DT among the earliest land plants, the occurrence of DT varies across extant land plant lineages and life stages. Many bryophytes have retained vegetative DT in the gametophyte stage (Oliver et al. 2005, and references therein). In extant vascular plants (tracheophytes), DT occurs in certain non-vegetative stages (e.g., seeds and pollen), but has largely been lost in vegetative tissues [with some notable exceptions, the so-called ‘resurrection plants’; Hartung et al. (1998); Kappen and Valladares (2007); Scott (2000)]. Instead, the majority of tracheophytes solely rely on avoiding desiccation with adaptations including a waxy cuticle, stomata, roots, and an extensive vascular system. Correlated with these differences in functional strategies to cope with water stress is a major difference in life cycle. Over time, plants shifted away from wholly gametophyte-dominated life cycle (as in extant bryophytes) to co-independence of the gametophyte and sporophyte (i.e., ferns and lycophytes) to complete loss of the independent gametophyte (i.e., spermatophytes). Little is known about the details of the transition from desiccation tolerance to avoidance, nor how this took place in conjunction with the switch from a gametophyte to a sporophyte-dominated life cycle; yet, these two steps underpin the radiation of vascular plants onto dry land (Puttick et al. 2018; Qiu et al. 2006; Wickett et al. 2014).

Ferns represent an important, understudied group of tracheophytes for investigating both the ecology and evolution of desiccation tolerance. Ferns are the only major lineage among land plants with sporophytes and gametophytes that are capable of growing independently from each other. These two life stages differ drastically in their morphology and physiology—fern sporophytes have true vascular tissue and are homoiohydric (i.e., they regulate their internal water content), whereas fern gametophytes are much smaller (often < 1 cm), lack vascular tissue, and are poikilohydric. Recent studies indicate that the gametophytes of many fern species are desiccation tolerant, while sporophytes of the same species are thought to lack DT (Pittermann et al. 2013; Watkins et al. 2007a). Thus, ferns present a unique opportunity to observe the transition from desiccation tolerance to desiccation avoidance across independently growing life stages within a single organism.

Filmy ferns (Hymenophyllaceae) represent a particularly interesting case for studying the ecological and evolutionary significance of vegetative DT across life stages in ferns. Filmy ferns are a large family [*c*. 430 spp.; Pteridophyte Phylogeny Group I (2016)] of primarily tropical, leptosporangiate ferns named for their remarkably thin leaf laminae, which are usually only a single cell layer in thickness and lack cuticle or stomata (Fig. 1a–d). Filmy fern sporophytes are a remarkable exception to the ‘typical’ homoiohydric fern sporophyte described above because they have reverted to poikilohydry: their thin laminae allow for passive, rapid uptake and loss of moisture, thereby maintaining nearly constant equilibrium with atmospheric humidity (Raven 1999). Many species of epiphytic filmy ferns have extremely reduced root systems (Schneider 2000), relying completely upon absorption through leaf laminae for water (Shreve 1911), although terrestrial species possess true roots and use ground water (Dubuisson et al. 2011). Thus, the morphological and physiological gap between sporophytes and gametophytes is arguably smaller in filmy ferns compared to any other fern clade. This makes filmy ferns a potentially useful group for understanding the transition from a desiccation tolerant, gametophyte-dominated strategy to a desiccation avoiding, sporophyte-dominated one, since the sporophytes and gametophytes of ancestral tracheophytes were also likely initially similar, then gradually diverged by a series of step-wise adaptations (Bateman et al. 1998; Ligrone et al. 2012).

**Fig. 1.**
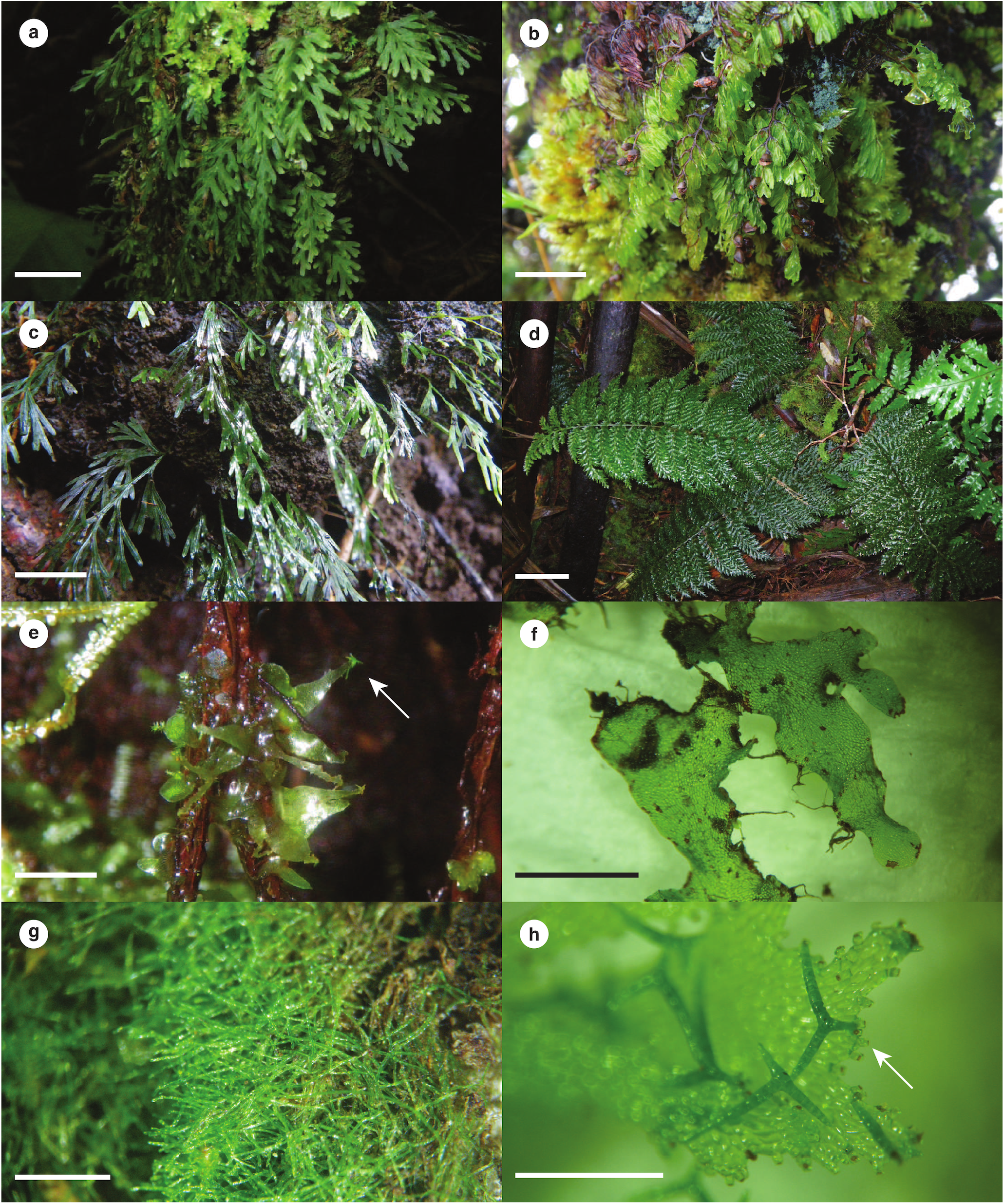
Examples of filmy ferns from French Polynesia. (**a**–**d**) sporophytes; (**e**–**h**) gametophytes. (**a**) *Crepidomanes humile*, a low elevation epiphyte. *Nitta 3392* (GH). (**b**) *Hymenophyllum multifidum*, a high elevation epiphyte growing in association with leafy liverworts and mosses. *Nitta 1257* (GH). (**c**) *Polyphlebium endlicherianum*, a saxicolous species found at low to middle elevations. *Nitta 629* (GH). (**d**) *Callistopteris apiifolia*, a terrestrial species with sporophytes occurring only in humid cloud forests. *Nitta 3990* (GH). (**e**) *Callistopteris apiifolia* (ribbon morphotype). Arrow indicates gemmae (asexual propagules) at tip of lobe. *Nitta 2565* (GH). (**f**) *Hymenophyllum polyanthos* (ribbon morphotype). *Nitta 3862* (GH). (**g**) Filamentous gametophytes of an unidentified species of Hymenophyllaceae. *Nitta 4036* (GH). (**h**) *Callistopteris apiifolia*, showing detail of gemmifers (attachment points of gemmae, example indicated by arrow) and branched gemmae. *Nitta 3887* (GH). Scalebars: (**a**–**c**) 1.0 cm; (**d**) 5.0 cm; (**e**) 5.0 mm; (**f**) 2.0 mm; (**g, h**) 1.0 mm. Photographs by J. H. Nitta.

Filmy fern sporophytes encompass a broad diversity of growth forms (terrestrial, saxicolous, hemi-epiphytic, and holoepiphytic) and morphologies, with rhizome and leaf sizes varying over an order of magnitude (Dubuisson et al. 2013; Hennequin et al. 2008; Nitta and Epps 2009). As would be expected from their poikilohydric nature, DT of varying degrees has been reported in sporophytes of many filmy fern species (see references below). Filmy fern gametophytes are similarly diverse, with morphotypes including filamentous, ribbon, and a combination of these two types (Fig. 1e–h). While there are no quantitative data about DT in filmy fern gametophytes available, DT has been reported in gametophytes from several other fern lineages (Pittermann et al. 2013; Watkins et al. 2007b; Watkins and Cardelús 2012), and may be expected to occur in filmy ferns as well. The possibility for varying degrees of vegetative DT in both stages of the life cycle of filmy ferns makes them a potentially useful system to compare how DT is utilized between haploid, non-vascular and diploid, vascular life stages. Furthermore, the diversity of growth forms and habitats in filmy ferns makes them an ideal group for a comparative study of water-stress strategies (i.e., desiccation tolerance vs. desiccation avoidance) and their association with different ecological niches (i.e., along gradients of temperature and relative humidity).

The degree of DT in filmy fern sporophytes occupying different niches has been relatively well studied. In a series of elegant experiments on Jamaican filmy ferns, Shreve (1911) demonstrated that water-use strategies differ between species with contrasting growth habits including terrestrial, low elevation epiphyte, and high elevation epiphyte. Proctor (2003, 2012) investigated water relations and the ability to recover from desiccation in sporophytes of several filmy ferns, and found a general correlation between high DT and adaptation for high light levels in widespread species vs. low DT and adaptation to low light levels in species occupying more sheltered habitats. In the only study to our knowledge including in situ physiological observations of hymenophyllaceous gametophytes, Johnson et al. (2000) found that gametophytes of *Vandenboschia speciosa* (Willd.) G. Kunkel (= *Trichomanes speciosum* Willd.) are adapted for extremely low light environments, and operate at lower light levels than conspecific sporophytes. Parra et al. (2009) and Saldaña et al. (2014) investigated the relationship between physiology (photosynthetic rates and water relations) and vertical stratification in several epiphytic filmy fern species (primarily genus *Hymenophyllum*) along tree trunks in Chile. They found that species occurring at greater heights in the trees tended to have greater DT, higher photosynthetic capacity (A_max_), and lower rates of evapotranspiration. A study using two species of *Hymenophyllum* demonstrated that DT in filmy ferns likely operates via a homoiochlorophyllous, constitutive mechanism, which is more similar to that of bryophytes than other drought-tolerant vascular plants, thus allowing filmy ferns to adjust to dry conditions extremely rapidly (Cea et al. 2014). This was further supported by evidence that induced biomechanical mechanisms likely do not play a role in DT of filmy ferns (Fallard et al. 2018; but see Mkhize et al. 2020). Recently, transcriptomics has been applied to elucidate the genes involved at each step of the desiccation-recovery cycle of filmy ferns (Ostria-Gallardo et al. 2020a, b). In epiphytic *Hymenoglossum cruentum* Cav., key genes and gene networks were identified related to intracellular mobility and photosynthetic metabolism in the desiccated state, and those related to detoxifying pathways and stabilization of photosystems during rehydration (Ostria-Gallardo et al. 2020b). In a comparison of low-trunk and canopy epiphytes, Ostria-Gallardo et al. (2020a) identified multiple differentially expressed genes that may be specific to tolerating desiccation in these microhabitats.

Despite this considerable interest in filmy fern ecophysiology, with the sole exception of Johnson et al. (2000), no studies have compared physiology between gametophyte and sporophyte stages, and we are aware of none that have done so across multiple species. Furthermore, to our knowledge, no physiological studies of filmy ferns have accounted for their phylogenetic relationships, and we lack an understanding of how physiological strategies have evolved in this clade. In addition to DT, photosynthetic optima (adaptation to different light levels) are likely to be important in determining the niches of life stages and species of filmy ferns (Flores-Bavestrello et al. 2016; Parra et al. 2015; Proctor 2003). Terrestrial and epiphytic species experience different levels of photosynthetic radiation and desiccation, and light intensity varies with height along the host plant in epiphytes (Benzing 1990; Watkins and Cardelús 2009). The combined effect of desiccation and light stress may induce greater oxidative damage in filmy ferns than either of these stressors alone (Niinemets et al. 2018), suggesting the need to integrate studies of DT with photosynthesis to gain a full picture of physiological adaptation in filmy ferns. Here, we leverage a recently developed DNA barcoding system (Nitta et al. 2017) to test several hypotheses related to DT and photosynthesis in a comparative phylogenetic framework including both filmy fern sporophytes and gametophytes: 1.) Light responses and DT differ between species with different growth habits. 2.) Light responses and DT differ between sporophytes and gametophytes. 3.) DT increases with decreasing environmental water availability. 4.) Gametophytes with broader elevational ranges have higher DT than those with with narrower ranges.

## Materials and methods

### Study system

We selected the island of Moorea, French Polynesia (17°32’ S, 149°50’ W) for our survey of filmy fern ecophysiology. Moorea is a small (135 km^2^; maximum elevation 1206 m) tropical oceanic island, located *c*. 15 km NW of its larger sister island, Tahiti. Despite its small size, it hosts a range of habitats including coastal strand, low elevation rainforest, and high elevation cloud forest. A total of 19 filmy fern species are known from Moorea, including multiple terrestrial, saxicolous, and epiphytic species (Florence in press; Murdock and Smith 2003; Nitta et al. 2011b, 2017; Ranker et al. 2005) (Figs 1, S1; all supplemental figures available in Online Resource 1). The two major clades of filmy ferns, the hymenophylloid and trichomanoid lineages (Pryer et al. 2001), are represented on Moorea by 7 and 12 species, respectively. We follow the taxonomic system of Ebihara et al. (2006) for Hymenophyllaceae.

### Field survey and DNA barcoding

We surveyed filmy ferns on Moorea as part of a larger study including all ferns of Moorea and Tahiti (Nitta et al. 2017). During this study, 17 sampling sites were established from *c*. 200 m to 1200 m primarily along trails to the summits of three large peaks on Moorea: Mt. Rotui (899 m), Mt. Mouaputa (880 m), and Mt. Tohiea (1206 m). At each site, sporophytes were sampled in a single 10 m × 10 m plot, and gametophytes sampled in two to three 50 cm × 50 cm plots located within the sporophyte plot. To estimate abundance of sporophytes, the 10 m × 10 m plot was subdivided into 25 2 m × 2 m subplots, and the presence of species in the sporophyte phase in each subplot was summed such that each species was assigned an abundance rank from 1 to 25. For the gametophyte plots, each 50 cm x 50 cm plot was subdivided into 25 5 x 5 cm squares, and a single gametophyte selected closest to the center of each square, to later be identified to species using DNA barcoding (see below). Epiphytes were sampled to a maximum height of *c*. 2 m on their host plants. For additional details on sampling procedure, see Nitta et al. (2017). The growth habit of each species was scored based on field observations as terrestrial, saxicolous, low elevation epiphyte (mostly occurring outside of cloud forest, i.e., below *c*. 500 m), or high elevation epiphyte (mostly occurring in cloud forest, i.e., above *c*. 500 m). *Crepidomanes bipunctatum* (Poir.) Copel. and *C. humile* (G. Forst.) Bosch were observed growing as both epiphytes and saxicoles, but were treated as epiphytes to distinguish them from exclusively saxicolous species.

Unlike sporophytes, fern gametophytes are cryptic and cannot be identified to species based on morphology alone. Therefore, we used DNA barcoding to identify filmy fern gametophytes to species following Nitta et al. (2017). Briefly, DNA extraction was performed using a modified CTAB protocol (Doyle and Doyle 1987) or the Plant Mini DNEasy kit (Qiagen, Valencia, California, USA) following the manufacturer’s protocol. Chloroplast *rbcL* and *trnH–psbA* were amplified and sequenced using primers and protocols following Nitta et al. (2017). For each marker, a reference library was constructed from sporophyte sequences using the ‘makeblastDB’ command in BLAST (Altschul et al. 1997). Field-collected gametophytes were identified by a local BLAST query against the reference library. Gametophytes matching > 99.5 % with a single reference sequence and no others were identified as that species; those that did not match any reference sequence or matched > 99.5 % with multiple reference sequences were not identified and excluded from further analysis. For this study, only gametophytes that could be identified as Hymenophyllaceae were used; others were excluded from analysis.

Fieldwork was done under permits issued by the French Polynesian Government (Délégation à la Recherche) and the Haut-commissariat de la République en Polynésie Francaise (Protocole d’Accueil 2012–2014). Voucher specimens were deposited at UC, with duplicates at GH and PAP (abbreviations follow Thiers (2021)).

### Microclimate

We used microclimate data collected by Nitta et al. (2020). This dataset includes relative humidity (RH) and temperature measured every 15 min from 2013-07-07 to 2014-07-05 at the fern survey sites with Hobo ProV2 dataloggers (Onset Computer Corporation, Bourne, Massachusetts, USA), and vapor pressure deficit (VPD) calculated from these two values. The dataloggers were outfitted with radiation shields (RS1 or RS3, Onset Computer Corporation, Bourne, Massachusetts, USA) to prevent direct contact with precipitation and solar radiation, and placed in pairs at each site (one ‘epiphytic’ datalogger at *c*. 1.5 m on a tree, one ‘terrestrial’ datalogger at *c*. 10 cm next to the tree). During the study period, some dataloggers malfunctioned, presumably due to prolonged exposure to high humidity. Any days during which one or more dataloggers failed are excluded from the dataset. One terrestrial datalogger (Mt. Tohiea 393 m site) was lacking data for more than half of the survey period, but had data available for the same period from a preliminary survey conducted during the previous year (2012–2013), so the previous year’s data for this site was used instead. The dataset includes 244 days of data from 15 sites (26 dataloggers). Mean daily temperature on Moorea is strongly correlated with elevation, dropping *c*. 6 °C from the lowest surveyed site (201 m) to the highest (1206 m) (*R*^2^ = 0.87; *P* < 0.001; linear model); temperature does not differ significantly between epiphytic and terrestrial habitats (Nitta et al. 2020). Epiphytic habitats tend to have lower daily mean RH, and greater daily variation in RH, than terrestrial ones. This difference is greatest at low elevations, and smallest at high elevations (Nitta et al. 2020).

### Desiccation tolerance

Samples were collected in the field and stored in plastic bags with a small amount of water to keep them fresh during transport to the lab. For sporophytes, eight individuals (one individual = single whole frond including a *c*. 2 cm section of rhizome) were used per treatment, and all individuals for each species came from a single population. We did not attempt to differentiate genotypes between fronds in species with long-creeping rhizomes, which often form clonal mats; hence, in some cases, individuals may be genetically identical ramets. All experiments were initiated within 48 h of collection. Pre-treatment maximum photochemical yield of photosystem II (F_v_/F_m_) (Kitajima and Butler 1975) was measured in fresh plants after a 10 min period of dark-adaptation using a portable mini-PAM chlorophyll fluorometer (Walz Gmbh, Effeltrich, Germany). Any samples with pre-treatment F_v_/F_m_ < 0.4 were excluded from analysis, as typical values of F_v_/F_m_ in healthy plants tend to be *c*. 0.8 (Björkman and Demmig 1987). Samples were then transferred to desiccation chambers containing saturated salts at three different desiccation intensities or a control treatment with moist tissues (100 % RH), and water withheld for either a short (2 d) or long (15 d) interval (Testo and Watkins 2013). Conditions inside the desiccation chambers were monitored during the experiment using Track-It RH/Temp dataloggers (Monarch Instrument, Amherst, NH) logging every 10 min (2012 field season for species other than *Callistopteris apiifolia* (Presl) Copel.) or Hobo ProV2 dataloggers logging every 5 min (2012 *C. apiifolia*, 2013, 2014 field seasons). The desiccation chambers were kept in an air-conditioned room. Temperature inside the chambers was mean 24.1 ± SD 1.15 °C (Fig. S2) (hereafter, the value following ‘±’ is standard deviation unless otherwise mentioned). Salts used for desiccation and their corresponding mean water potentials and approximate relative humidity and VPD are as follows: LiCl (−282 MPa, 18 % RH, 2.45 kPa), Mg(NO_3_)_2_ (−86 MPa, 58 % RH, 1.25 kPa), and NaCl (−38 MPa, 80 % RH, 0.60 kPa). Following the desiccation treatment, plants were rewetted by wrapping them in moist tissues using water from a local stream. Plants were allowed to rehydrate, and F_v_/F_m_ was again measured at 0.5 h, 24 h, and 48 h following rewetting. Ability to recover from desiccation was quantified by comparing pre- and post-treatment F_v_/F_m_ (Watkins et al. 2007a).

We measured relative water content (RWC) in a subset of samples by recording the mass of samples prior to the DT test (fresh mass), at each step of the DT test (turgid mass), then following the DT test after drying them overnight in an oven at 65 °C (dry mass). Relative water content was calculated as 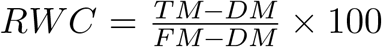, where *TM* is turgid mass, *DM* is dry mass, and *FM* is fresh mass.

Because field-collected gametophytes could not be identified to species prior to DNA extraction, no planned replication by species was possible. We therefore applied the same treatment for all gametophytes (2 d at -86 MPa) in batches of individuals (Table S1), and verified the species of each individual after the experiment using the DNA barcode approach described above. Thus, gametophytes of some species were sampled many times whereas those of other species were not. Relative water content was not calculated for gametophytes, as these were too small to measure accurately with the balances available.

### Light responses

Rapid light response curves were constructed for each species by measuring photosynthetic yield at gradually increasing levels of photosynthetically active actinic light (400 nm to 700 nm) with the Light Curve function of the mini-PAM portable chlorophyll fluorometer as described by the manufacturer. Briefly, the maximum yield (F_m_) was first measured in the absence of actinic light. Next, plants were equilibrated to the actinic light for 30 s, and photosynthetic yield (ΔF/F_m_) measured with a 0.8 s saturating pulse. This was repeated for each photosynthetic photon flux density (PPFD) level, up to ca. 500 µmol·m^-2^·s^-1^. Response curves were fitted using the equation *y* = *A*(1 − *e*^−*kx*^), where *y* is relative electron transport rate of photosystem II (RETR), *x* is photosynthetic photon flux density (PPFD), *A* is the asymptote of the curve, and *k* is a slope parameter; outliers at high PPFD likely to distort the curve were discarded (Proctor 2012). The PPFD at 95 % saturation of RETR (PPFD_95%_) and maximum RETR (ETR_max_) were then calculated from the curve. Light responses were measured in eight individuals per species in the lab for sporophytes, and on single individuals in the field for gametophytes. The gametophytes were later identified to species using the DNA barcodes.

### Statistical analysis

All statistical analyses were done in R v.4.0.3 (R Core Team 2020).

We derived the phylogenetic tree used in this study by extracting Hymenophyllaceae from the tree of Nitta et al. (2017) with the ‘extract.clade’ function of the ‘ape’ package (Paradis et al. 2004). This tree is in agreement with previously published, more densely sampled phylogenies (Dubuisson et al. 2003; Ebihara et al. 2007; Hennequin et al. 2006; Pryer et al. 2001).

We quantified the degree of phylogenetic signal in physiological traits using Blomberg’s *K* (Blomberg et al. 2003) and Pagel’s λ (Freckleton et al. 2002; Pagel 1999) as implemented with the ‘phylosig’ function in the ‘phytools’ package (Revell 2012). Both measures test the hypothesis that the trait of interest is evolving according to Brownian Motion (BM). λ is a scaling parameter that ranges from zero to one: λ near zero indicates random distribution of trait values across the tree, and λ near one indicates evolution of traits along the phylogeny following BM. For *K*, values near one indicate evolution of traits following BM; *K* > 1 indicates traits more conserved than expected under BM, and *K* < 1 indicates that traits have less phylogenetic signal than expected under BM. We tested the significance of *K* by comparing the observed value against values from a null distribution of 10,000 phylogenies with the traits randomly shuffled across the tips. We tested the significance of λ with a log-likelihood test comparing the likelihood of the observed value of λ vs. λ = 0. For all further analyses, we used methods that account for phylogenetic history for physiological traits that showed evidence of phylogenetic signal in at least one of the two metrics, and standard methods otherwise.

We used general linear mixed models (GLMMs) to investigate the effect of growth habit and life stage on DT (% recovery of F_v_/F_m_ after 48 h recovery from desiccation at -86 MPa for 2 d) and photosynthetic parameters (ETR_max_ and PPFD_95%_) using the ‘MCMCglmm’ package (Hadfield 2010). For each of the three response variables (species mean DT, ETR_max_, and PPFD_95%_), we constructed five models: a null model with no fixed effects, or models with either growth habit (high-elevation epiphyte, low-elevation epiphyte, terrestrial or saxicolous), life stage (sporophyte vs. gametophyte), growth habit and life stage, or growth habit, life stage, and their interaction as fixed effects. We included species as a random effect in all models, and included a phylogenetic variance-covariance matrix if the response variable showed evidence of phylogenetic signal (Hadfield 2010). We used an inverse-Gamma distribution for all priors, and ran analyses for 500,000 iterations, with burnin after 1,000 iterations and thinning every 500 iterations. We compared models using the Deviance Information Criterion (DIC) (Spiegelhalter et al. 2002) and selected the best-fitting model as the one with the lowest DIC.

Comparisons of DT, ETR_max_, and PPFD_95%_ between gametophytes and sporophytes of the same species do not involve phylogeny, so we used a two-sided *t*-test for these.

To test for a correlation between environmental water availability and DT, we first derived a single maximum VPD for each species by calculating the average of the maximum daily VPD at all sites where that species was present. We used maximum VPD (i.e., minimum humidity) values because these may represent a climatic extreme that limits the occurrence of species. We used data from the ‘epiphytic’ dataloggers mounted at 1.5 m on trees for epiphytic species and data from the ‘terrestrial’ dataloggers on the ground for non-epiphytic species. Data from the Mt. Rotui 830 m site were excluded because this site is much more exposed than the other high elevation sites, and appears as a significant outlier in the microclimatic data (Nitta et al. 2020). Since DT showed significant phylogenetic signal (see Results), we used phylogenetic generalized least squares (PGLS) (Freckleton et al. 2002) to test for a relationship between VPD and DT while taking phylogeny into account, using species’ means.

To test whether gametophytes with broader ranges have higher DT, we calculated the extent of gametophyte range beyond sporophytes (“G beyond S”) as the total elevational range (m) gametophytes were observed growing beyond sporophytes at either end of the elevational gradient. We conducted PGLS with DT as the response variable and G beyond S as the explanatory variable with the ‘pgls’ function in the ‘caper’ (Orme et al. 2018).

Datasets used in this study are available on FigShare (https://doi.org/10.6084/m9.figshare.14184572). Code used to analyze the data and generate figures and this manuscript are available on GitHub (https://github.com/joelnitta/moorea_filmies). A Docker image is available to run the code at https://hub.docker.com/r/joelnitta/moorea_filmies. The ‘targets’ package (Landau 2021) was used to automate the workflow. The ‘ggplot2’ (Wickham 2016), ‘ggtree’, (Yu et al. 2017) and ‘patchwork’ (https://github.com/thomasp85/patchwork) packages were used to generate the figures.

*** NOTE TO READER: The DOI for the dataset will not be active until the paper is accepted. Until then, this link can be used for the dataset https://figshare.com/s/d6349abf01a3756a5aae ***

## Results

### Field survey

In total, 22 filmy fern taxa (including two species that were recognized as multiple varieties) were observed on Moorea. Three taxa with distinct *rbcL* sequences, morphology, and elevational ranges could be distinguished within the *Crepidomanes minutum* (Blume) K. Iwats. species complex, but species status of these taxa remains uncertain (Nitta et al. 2011a). We refer to these here as *C. minutum* var. 1, 2, and 3. Similarly, two members of the *Abrodictyum asae-grayi* (Bosch) Ebihara & K. Iwats. complex (also including *Abrodictyum caudatum* (Brack.) Ebihara & K. Iwats. in our tree; Fig. S1) were observed on the basis of *rbcL* and morphology, but as taxonomic treatment is beyond the scope of the current study we refer to them as *A. asae-grayi* var. 1 and 2.

In several species, gametophytes exceeded the maximum or minimum elevational range of sporophytes including *C. apiifolia, C. minutum* var. 2 and 3, *Hymenophyllum digitatum* (Sw.) Fosberg, *Hymenophyllum javanicum* Spreng., *Hymenophyllum pallidum* (Blume) Ebihara & K. Iwats., and *Polyphlebium borbonicum* (Bosch) Ebihara & Dubuisson (Fig. S3). All of these have sporophytes that are normally restricted to cloud forest (*c*. 500–1200 m), but gametophytes that are distributed to lower elevations.

Sporophytes of two taxa, *C. minutum* var. 3 and *Hymenophyllum braithwaitei* Ebihara & K. Iwats., did not occur in the study plots, but each was observed from a single population on Mt. Tohiea, at *c*. 1000 m and near the summit (1206 m), respectively. We include these observations for comparison of ranges between sporophytes and gametophytes.

### Phylogenetic signal

For DT, *K* was near 1 with *P* < 0.1 in both life stages (gametophytes *K* = 0.68, *P* = 0.036; sporophytes *K* = 0.69, *P* = 0.053); λ was near 1 for both life stages, but not significant for sporophytes (gametophytes λ = 0.68, *P* = 0.063; sporophytes λ = 0.71, *P* = 0.298). There was no evidence of significant phylogenetic signal in photosynthetic parameters (PPFD_95%_ and ETR_max_) in either life stage (*P* » 0.05, Table 1). Therefore, phylogenetic methods were used for analyses involving DT, but not photosynthetic parameters.

**Table 1.**
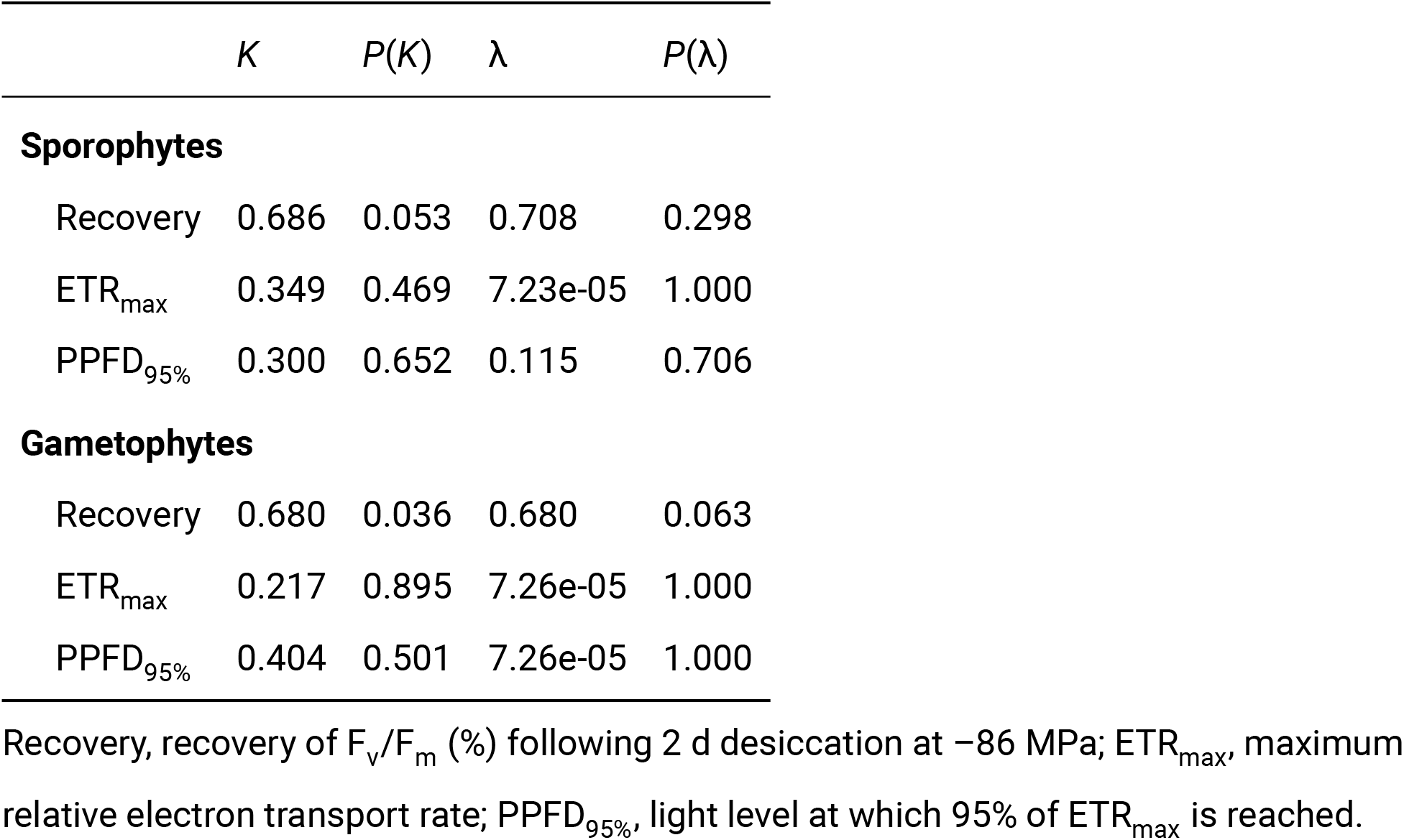
Results of tests for phylogenetic signal (Blomberg’s *K* and Pagel’s λ) in desiccation tolerance (DT) and photosynthetic parameters of filmy ferns from Moorea, French Polynesia by life stage

| | $K$ | $P(K)$ | $\lambda$ | $P(\lambda)$ |
| --- | --- | --- | --- | --- |
| **Sporophytes** |  |  |  |  |
| Recovery | 0.686 | 0.053 | 0.708 | 0.298 |
| ETR<sub>max</sub> | 0.349 | 0.469 | 7.23e-05 | 1.000 |
| PPFD<sub>95%</sub> | 0.300 | 0.652 | 0.115 | 0.706 |
| **Gametophytes** |  |  |  |  |
| Recovery | 0.680 | 0.036 | 0.680 | 0.063 |
| ETR<sub>max</sub> | 0.217 | 0.895 | 7.26e-05 | 1.000 |
| PPFD<sub>95%</sub> | 0.404 | 0.501 | 7.26e-05 | 1.000 |

Recovery, recovery of $F_v/F_m$ (%) following 2 d desiccation at $-86$ MPa; ETR<sub>max</sub>, maximum relative electron transport rate; PPFD<sub>95%</sub>, light level at which 95% of ETR<sub>max</sub> is reached.

### Desiccation tolerance

Ability to recover from desiccation was tested in sporophytes of 14 species and gametophytes of 14 species, including 12 species with both life stages.

A general correlation of DT with habitat was observed in sporophytes (Fig. 2a). The two terrestrial species lacked DT, although *C. apiifolia* showed slightly greater ability to recover by 48 h compared to *Abrodictyum dentatum* (Bosch) Ebihara & K. Iwats., which completely failed to recover from even the gentlest treatment of 2 d at –38 NaCl (more stringent tests were not conducted on *A. dentatum* after confirming its lack of DT at this level). Saxicolous species (*Crepidomanes kurzii* (Bedd.) Tagawa & K. Iwats., *Polyphlebium endlicherianum* (C. Presl) Ebihara & K. Iwats., and *Vandenboschia maxima* (Blume) Copel.) also had low DT; only *C. kurzii* was able to recover from desiccation up to –86 MPa for 2 d but not more. Nearly all epiphytic species were capable of recovering from desiccation of up to 2 d at –86 MPa. *Crepidomanes bipunctatum*, a widespread low-elevation epiphyte, showed the greatest ability to recover from desiccation across all treatments. Interestingly, high-elevation epiphytes *Hymenophyllum multifidum* (G. Forst.) Sw. and *Hymenophyllum polyanthos* (Sw.) Sw. could recover from desiccation at –86 or –282 MPa for 15 d, but not –38 MPa.

**Fig. 2.**
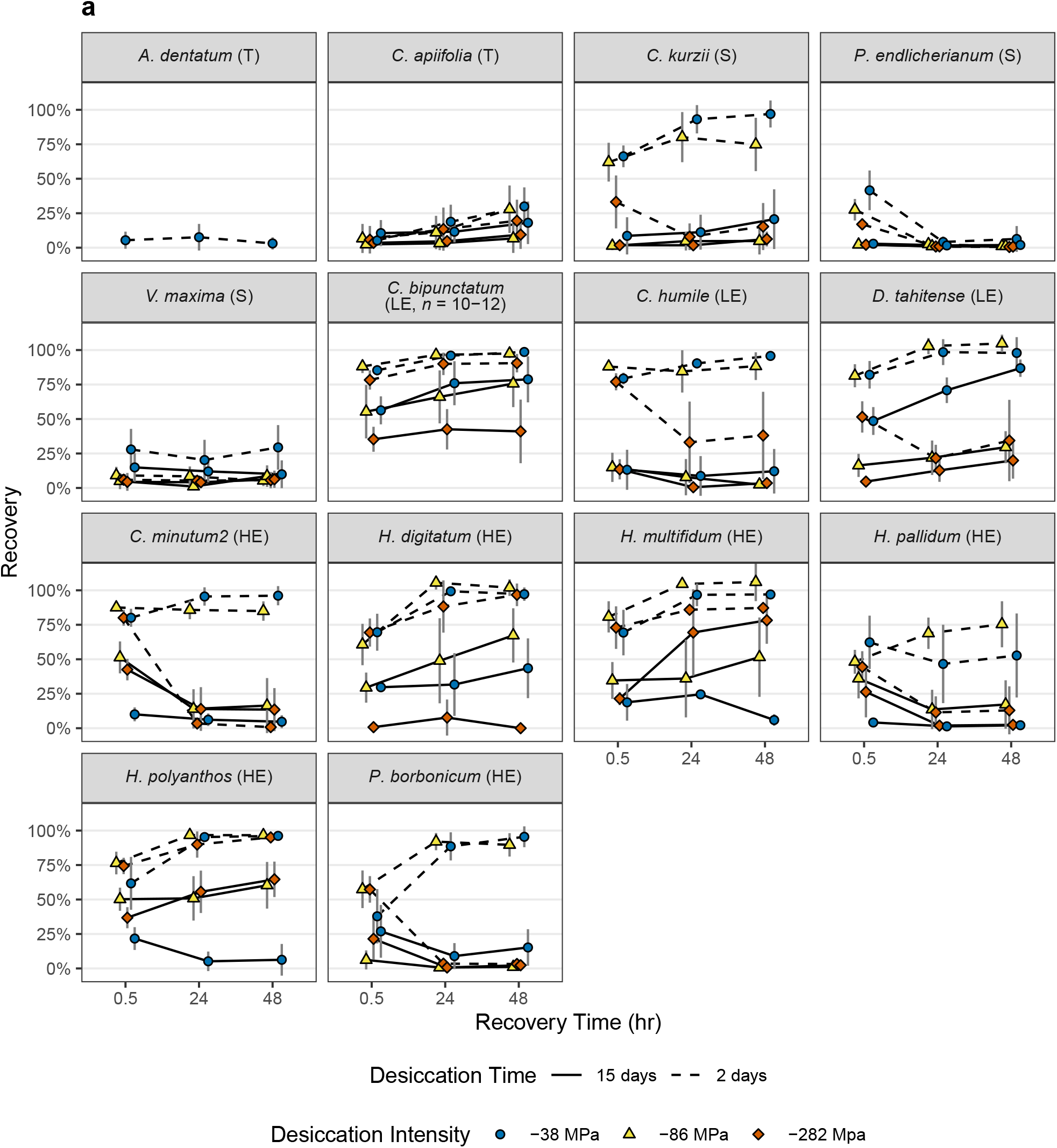
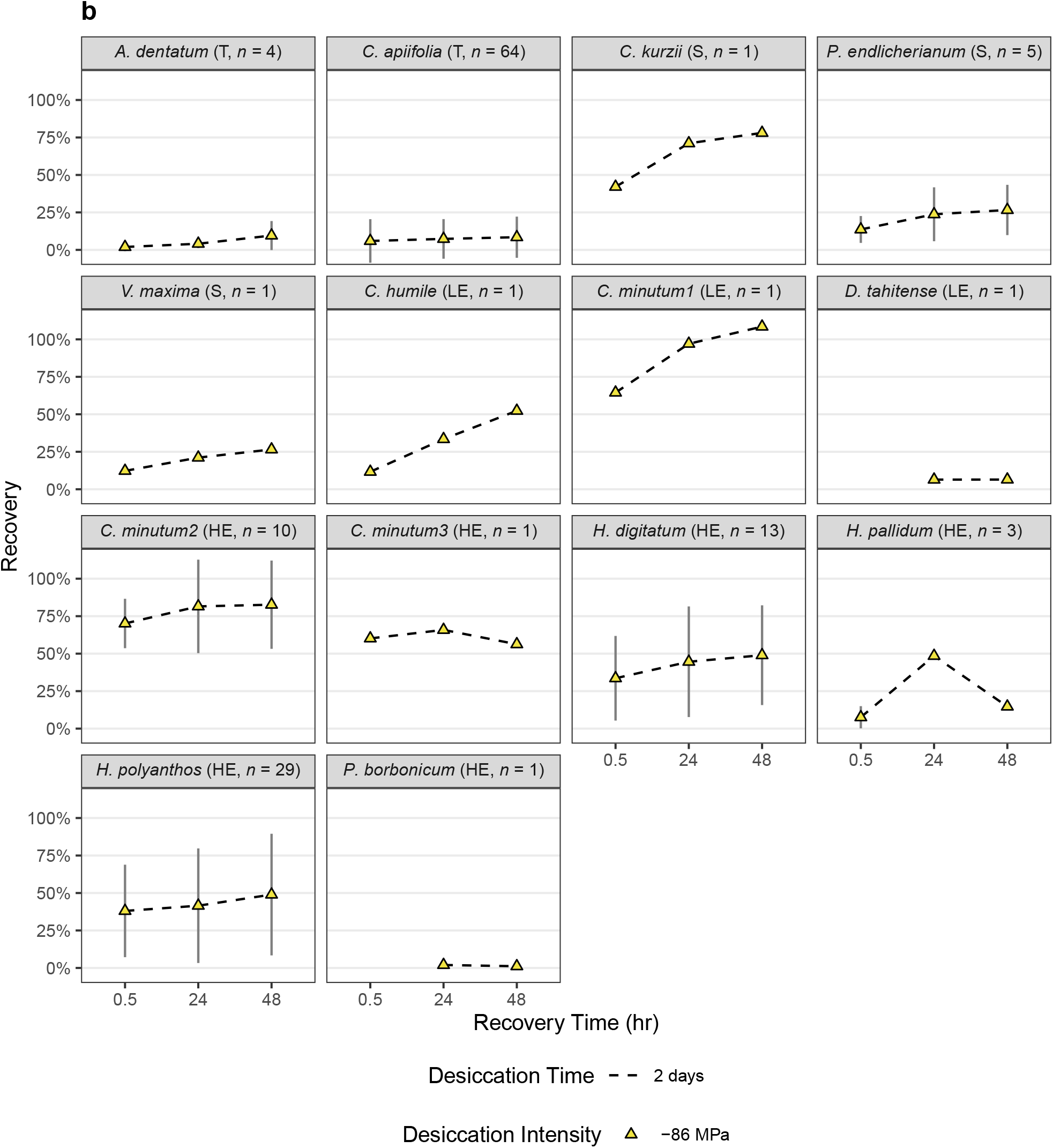
Desiccation tolerance of filmy fern sporophytes (**a**) and gametophytes (**b**) from Moorea, French Polynesia. Recovery was measured by comparing maximum photochemical yield of photosystem II before desiccation treatment with values at 30 min, 24 hr, and 48 hr following desiccation at three different intensities for either 2 or 15 days (see Methods). Recovery was not measured for all combinations of species and desiccation treatments; missing points indicate data not collected (not recovery of 0%). Error bars indicate standard deviation. *n* = 8 samples per species/treatment unless otherwise indicated. Growth habit indicated in parentheses after each species: T, terrestrial; S, saxicolous; LE, low elevation epiphyte; HE, high elevation epiphyte

Gametophytes of terrestrial species could not tolerate desiccation (mean 48 h recovery 9.0 % ± 0.8 %, *n* = 2 species). Levels of DT were in general higher for saxicolous and epiphytic gametophytes, but otherwise did not show a clear pattern (Fig. 2b).

Desiccation tolerance of gametophytes tended to be lower than sporophytes (Fig. 3), and was significantly lower in *C. apiifolia, H. polyanthos, H. pallidum*, and *H. digitatum* (*t*-test, *P* = 0.015, < 0.001, 0.035, and < 0.001, respectively). Only *P. endlicherianum* showed significantly higher DT in gametophytes compared to sporophytes (*t*-test, *P* = 0.027).

**Fig. 3.**
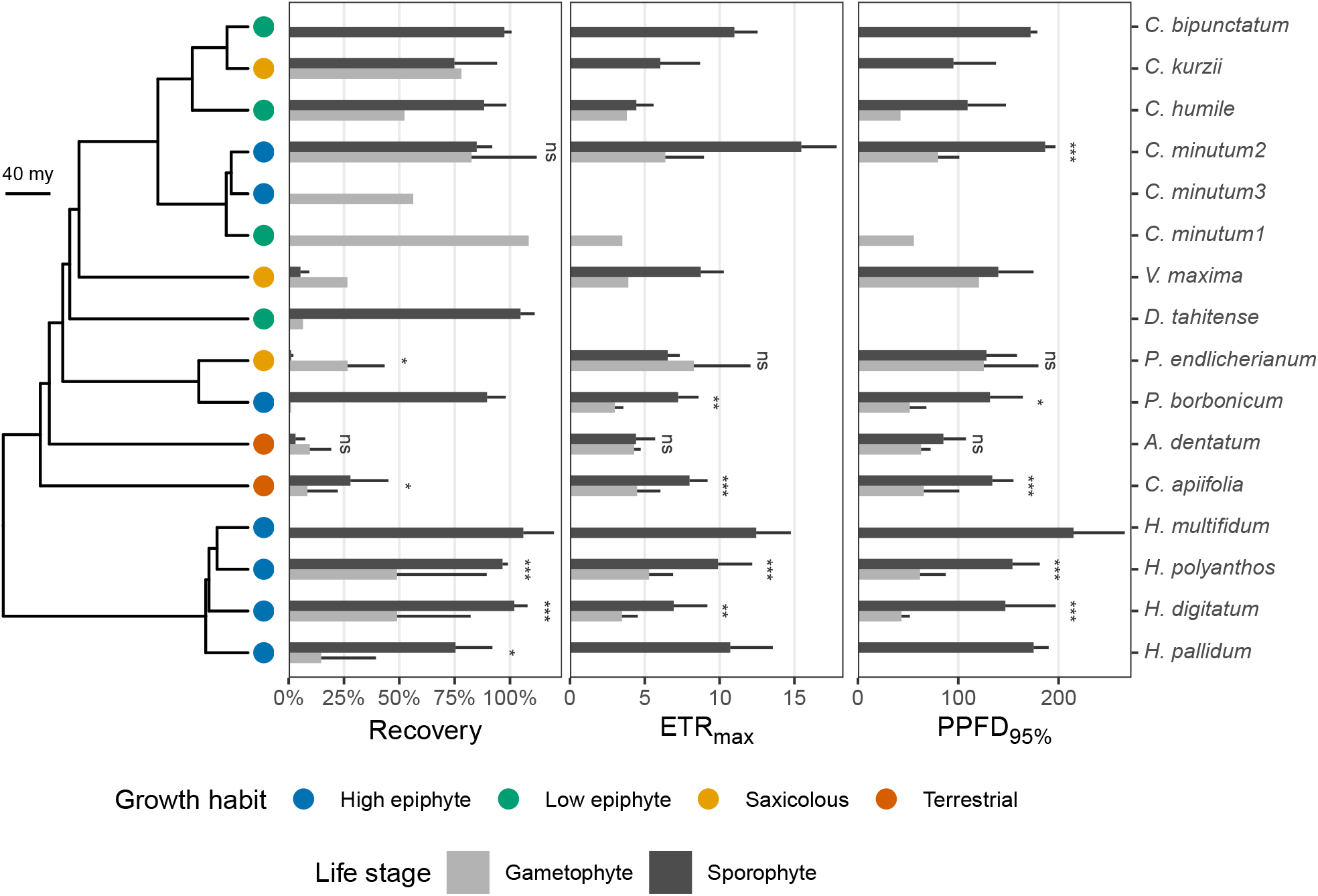
Phylogeny of filmy ferns from Moorea, French Polynesia with comparison of physiological parameters (mean ± 1 SD) between sporophytes (dark grey bars) and gametophytes (light grey bars). Phylogeny adapted from Nitta et al. (2016); only species with physiological data shown. Bootstrap support >50% from 100 ML bootstraps indicated at nodes. Bold letters indicate major clades (H = hymenophylloid clade, T = trichomanoid clade). Growth habit indicated by colored symbols (blue = low elevation epiphyte, green = high elevation epiphyte, yellow = saxicolous, red = terrestrial). *Crepidomanes bipunctatum* and *C. humile* were observed growing as both epiphytes and saxicoles, but are shown as epiphytes to distinguish them from exclusively saxicolous species. Asterisk indicates significant difference in means (*P* <0.05) between life stages for a particular species (*t*-test); ns indicates no significant difference; lack of ns or asterisk means not enough observations available for that species for *t*-test. Left bar plot shows recovery (%) of chlorophyll fluorescence (F_v_/F_m_) following 2 day desiccation at –86 MPa (*Abrodictyum dentatum* was desiccated at –38 MPa instead of –86 MPa). Middle bar plot shows maximum relative electron transport rate of photosystem II (ETR_max_) (µmol e-·m^-2^·s^-1^). Right bar plot shows photosynthetic photon flux density at 95% saturation of RETR (PPFD_95%_) (µmol quanta·m^-2^·s^-1^). Absence of a bar indicates measurements not available for gametophytes or sporophytes of that species.

Control samples generally did not show any decrease in F_v_/F_m_ throughout the duration of the experiment (Fig. S4) and maintained RWC near 100% (Figs S5, S6). Relative water content in the desiccated state ranged from 5.1 % ± 5.2 % (*n* = 9 species) in the –282 MPa, 15 d treatment to 17.2 % ± 8.0 % (*n* = 10 species) in the –38 MPa, 2 d treatment (Fig. S5), and returned to *c*. 100 % upon re-wetting in all species sampled (Figs S5, S6).

Relative humidity in the growth chambers was high when samples were first inserted, then gradually decreased to the target value maintained by each drying salt (Fig. S2). In a few cases (particularly species with larger fronds, e.g., *C. apiifolia*), the NaCl chamber did not reach equilibrium humidity before 2 d (Fig. S2). Brief fluctuations in humidity were observed when chambers were opened to insert or remove samples (Fig. S2).

The GLMM including the combined effects of life stage plus growth habit and their interaction was selected as the best-fitting model for DT (Table 2). Sporophytes had 52.3% greater DT (95 % CI 24.1% to 77.1%) relative to gametophytes (Table 3).

**Table 2.** Deviance information criterion (DIC) and delta DIC for general linear mixed models (GLMMs) of photosynthetic parameters and desiccation tolerance in response to life stage and growth habit in filmy ferns from Moorea, French Polynesia

| Fixed effects | DIC | delta DIC |
| --- | --- | --- |
| **ETR<sub>max</sub>** |  |  |
| **Life stage** | **112.39** | **0.00** |
| Life stage + habit | 114.52 | 2.13 |
| Life stage + habit + Life stage:habit | 115.69 | 3.30 |
| Null | 122.68 | 10.29 |
| Habit | 125.55 | 13.16 |
| **PPFD<sub>95%</sub>** |  |  |
| **Life stage + habit + Life stage:habit** | **211.05** | **0.00** |
| Life stage | 230.46 | 19.42 |
| Life stage + habit | 232.97 | 21.93 |
| Null | 248.08 | 37.04 |
| Habit | 251.78 | 40.73 |
| **Recovery** |  |  |
| **Life stage + habit + Life stage:habit** | **8.01** | **0.00** |
| Life stage | 16.36 | 8.35 |
| Life stage + habit | 16.77 | 8.76 |
| Habit | 23.28 | 15.27 |
| Null | 23.39 | 15.38 |

ETR<sub>max</sub>, maximum relative electron transport rate; PPFD<sub>95%</sub>, light level at which 95% of ETR<sub>max</sub> is reached; Recovery, recovery of $F_v/F_m$ (%) following 2 d desiccation at –86 MPa
Best model (that with the lowest DIC) for each response variable in bold

**Table 3.** Predictor effects, lower and upper 95% confidence interval (CI), effective sample size, and significance of model parameters for the best general linear mixed models (GLMMs) of photosynthetic parameters and desiccation tolerance in response to life stage and growth habit in filmy ferns from Moorea, French Polynesia

| Fixed effects | Predictor | Mean effect | Lower 95% CI | Upper 95% CI | Ef. sample size | P MCMC |
| --- | --- | --- | --- | --- | --- | --- |
| **ETR<sub>max</sub>** |  |  |  |  |  |  |
| **Life stage** | **Sporophyte** | **3.880** | **1.595** | **6.180** | **997** | **0.002** |
| **PPFD<sub>95%</sub>** |  |  |  |  |  |  |
| Life stage + habit + Life stage:habit | Low Epiphyte | -9.964 | -53.576 | 35.593 | 998 | 0.653 |
| **Life stage + habit + Life stage:habit** | **Saxicolous** | **55.087** | **5.859** | **106.377** | **997** | **0.034** |
| Life stage + habit + Life stage:habit | Terrestrial | 0.133 | -44.127 | 49.057 | 997 | 0.998 |
| **Life stage + habit + Life stage:habit** | **Sporophyte** | **105.200** | **77.411** | **135.288** | **998** | **0.001** |
| Life stage + habit + Life stage:habit | Low Epiphyte:Sporophyte | -18.553 | -77.984 | 34.518 | 998 | 0.447 |
| **Life stage + habit + Life stage:habit** | **Saxicolous:Sporophyte** | **-103.496** | **-156.991** | **-52.861** | **998** | **0.001** |
| **Life stage + habit + Life stage:habit** | **Terrestrial:Sporophyte** | **-59.072** | **-117.259** | **-11.485** | **998** | **0.040** |
| **Recovery** |  |  |  |  |  |  |
| Life stage + habit + Life stage:habit | Low Epiphyte | 0.122 | -0.268 | 0.545 | 997 | 0.513 |
| Life stage + habit + Life stage:habit | Saxicolous | 0.119 | -0.309 | 0.600 | 998 | 0.589 |
| Life stage + habit + Life stage:habit | Terrestrial | -0.239 | -0.793 | 0.371 | 1245 | 0.383 |
| **Life stage + habit + Life stage:habit** | **Sporophyte** | **0.523** | **0.241** | **0.771** | **997** | **0.001** |
| Life stage + habit + Life stage:habit | Low Epiphyte:Sporophyte | -0.074 | -0.537 | 0.405 | 998 | 0.745 |
| **Life stage + habit + Life stage:habit** | **Saxicolous:Sporophyte** | **-0.683** | **-1.130** | **-0.234** | **998** | **0.008** |
| Life stage + habit + Life stage:habit | Terrestrial:Sporophyte | -0.461 | -0.975 | 0.042 | 998 | 0.072 |

ETR<sub>max</sub>, maximum relative electron transport rate; PPFD<sub>95%</sub>, light level at which 95% of ETR<sub>max</sub> is reached; Recovery, recovery of F<sub>v</sub>/F<sub>m</sub> (%) following 2 d desiccation at -86 MPa Effect of life stage is relative to gametophytes; effect of growth habit is relative to high elevation epiphytes
Significant predictors in bold

### Photosynthetic optima

Light responses were measured in sporophytes of 13 species and gametophytes of 10 species, including nine species with both life stages.

Photosynthetic optima showed clear differences across life stages: sporophytes were adapted for higher light levels, whereas gametophytes were adapted for lower light levels (Figs 3, S7, S8). Sporophytes reached their maximum photosynthetic rate (PPFD_95%_) at light levels ca. 144.1 ± 36.8 µmol·m^−2^ ·s^−1^ (*n* = 13 species), whereas gametophytes did so at 71.0 ± 29.6 µmol·m^−2^ ·s^−1^ (*n* = 10 species). Similarly, sporophytes attained higher rates of electron transport (ETR_max_) (8.6 ± 3.2 µmol·m^−2^ ·s^−1^, *n* = 13 species) than gametophytes (4.6 ± 1.6 µmol·m^−2^ ·s^−1^, *n* = 10 species).

The GLMM including the effect of life stage only was selected as the best-fitting model for ETR_max_, and the combined effect of growth habit plus life stage and their interaction was selected for PPFD_95%_ (Table 2).

### Relationships of vapor pressure deficit and range size with DT

Desiccation tolerance increased significantly with VPD for sporophytes, but not gametophytes (PGLS, *P* = 0.038; Fig. 4; Table 4).

**Fig. 4.**
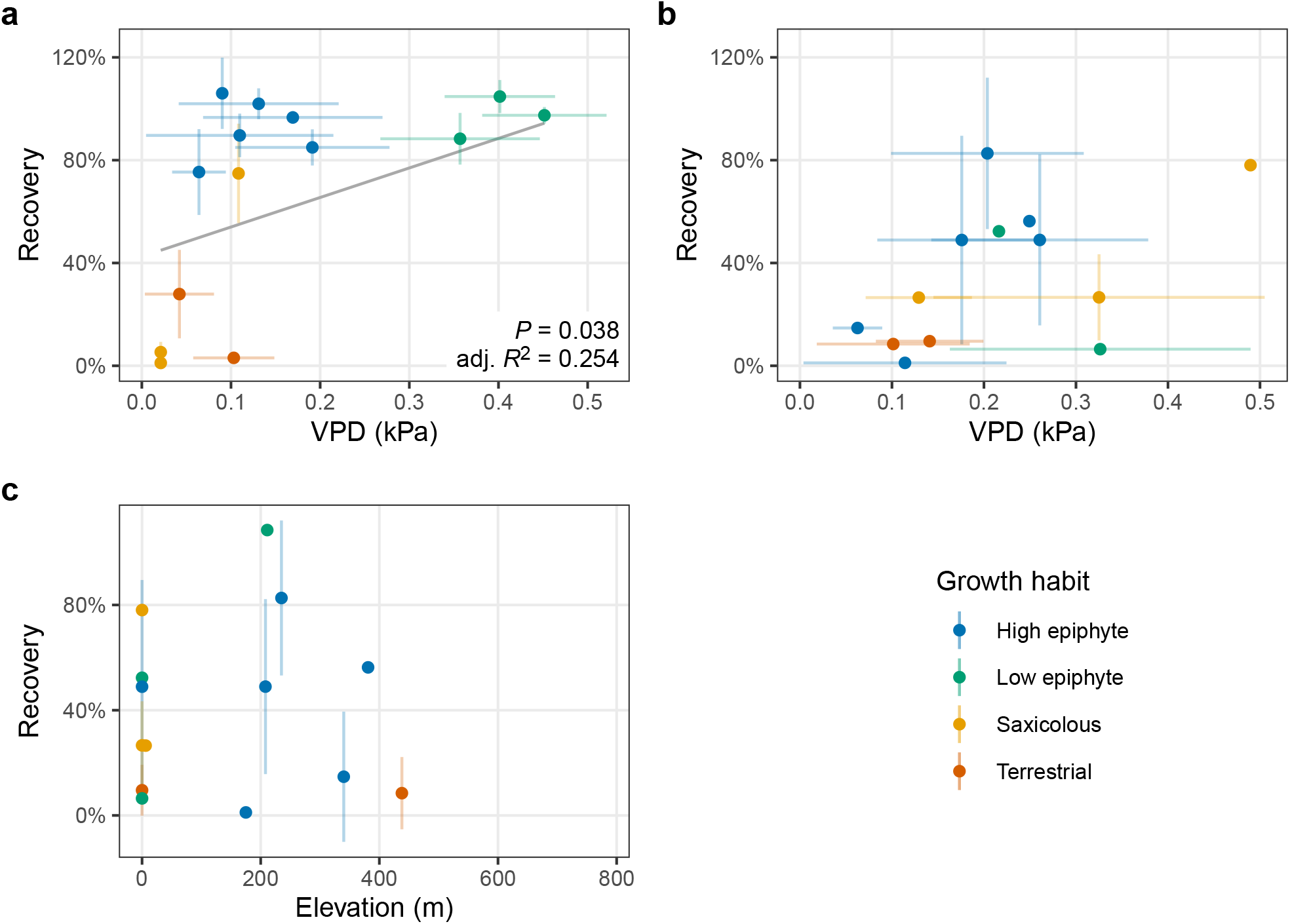
Recovery of F_v_/F_m_ (%) following 2 d desiccation at –86 MPa plotted against environmental vapor pressure deficit (VPD; kPa) (**a, b**) or range of gametophytes beyond sporophytes (m) (**c**) for gametophytes (**a, c**) or sporophytes (**b**). Species means ± 1 SD are shown. Growth habit indicated by colored symbols (blue = low elevation epiphyte, green = high elevation epiphyte, yellow = saxicolous, red = terrestrial). *Crepidomanes bipunctatum* and *C. humile* were observed growing as both epiphytes and saxicoles, but were treated as epiphytes to distinguish them from exclusively saxicolous species. *Abrodictyum dentatum* sporophytes were desiccated at –38 MPa instead of –86 MPa. Range of gametophytes beyond sporophytes calculated as the total elevation that gametophytes exceeded sporophytes at either end of the elevational gradient. Relationships between recovery and VPD or range of gametophytes beyond sporophytes were analyzed with phylogenetic generalized least squares (PGLS); significant relationships (*P* < 0.05) shown with grey line.

**Table 4.** Phylogenetic generalized least squares (PLGS) models of desiccation tolerance as a function of vapor pressure deficit (VPD) or the range of gametophytes beyond sporophytes (G beyond S) in filmy ferns from Moorea, French Polynesia

| Life stage | Predictor | df | *F* | adj. $R^2$ | *P* |
| --- | --- | --- | --- | --- | --- |
| S | VPD | 2,12 | 5.421 | 0.254 | 0.038 |
| G | VPD | 2,11 | 3.586 | 0.177 | 0.085 |
| G | G beyond S | 2,12 | 4.127 | 0.194 | 0.065 |

S, sporophyte; G, gametophyte

No significant relationship was detected between range size of gametophytes beyond sporophytes and DT (Fig. 4, Table 4).

## Discussion

Vegetative DT was a key innovation linked to the rise of early land plants, but is largely limited to non-vascular groups such as bryophytes and algae among extant plants. Filmy ferns have received considerable attention for their remarkable reversion to vegetative DT in the sporophyte stage (e.g., Cea et al. 2014; Ostria-Gallardo et al. 2020a; Proctor 2003; Shreve 1911). Yet, no previous study to our knowledge has evaluated parallel physiology of sporophytes and gametophytes of filmy ferns in relation to DT. Here, we present the first comparisons of physiological parameters between life stages within filmy ferns in a phylogenetic comparative framework.

### Filmy fern sporophytes have a wide range of DT

We identified a wide variation of DT in filmy fern sporophytes on Moorea, from marked sensitivity in terrestrial species to tolerance of water potentials well below -200 MPa (VPD > 2.45 kPa) in some epiphytes. Levels of similarly extreme DT have also been reported in other species of *Hymenophyllum* (Proctor 2003, 2012) and a general correlation of habitat with DT in filmy ferns has been previously established (Parra et al. 2009; Proctor 2003, 2012; Saldaña et al. 2014; Shreve 1911), but no other studies to our knowledge have documented a similar range of DT values in filmy ferns from a single site. One may wonder then, what is the adaptive significance of being able to withstand drying at -282 MPa (*c*. 2.45 kPa) vs. -86 MPa (*c*. 1.25 kPa) when the ambient VPD is typically < *c*. 0.10 kPa and only rarely exceeds > 0.50 kPa. The recent finding of a constitutive mechanism for DT in filmy ferns (Cea et al. 2014) supports the need for a relatively wide “safety margin,” and similar (or even more extreme) levels of constitutive DT are well known in bryophytes (Oliver et al. 1993). The conservative behavior of fern stomata, which lack a response to the plant stress hormone ABA, is well documented (Brodribb and McAdam 2011; McAdam and Brodribb 2012, 2013). It is possible that filmy ferns also behave conservatively with respect to water-stress. We suggest that the variation in DT observed here is relevant because it is correlated with habitat, and likely reflects overall physiological tolerance, not just artificial extremes. For example, sporophytes of some high elevation epiphytes that we surveyed (*H. multifidum, H. polyanthos*) seem to be able to better withstand intense vs. moderate desiccation, with higher recovery values at -282 and -86 MPa vs. -38 MPa when dry for 15 d. This may be an adaptation to cloud forest conditions; although cloud forests are frequently moist due to constant fog, dry periods are intense during the times when there is no cloud cover (J. Nitta, pers. obs.). The complete lack of DT in terrestrial species is not surprising, considering they have much more direct access to water than epiphytes and therefore are under substantially less selective pressure to maintain DT. Because controls kept at 100% RH maintained F_v_/F_m_ values near pre-treatment levels in all sporophytes (Fig. S4), it is unlikely that the lack of tolerance seen in terrestrial species is solely due to removal from the soil. The significant relationship between average daily maximum VPD and DT in sporophytes supports the specialization of sporophyte DT according to niche, and indicates that sporophytes of non-tolerant terrestrial and saxicolous species may be unable to occur in habitats with VPD exceeding *c*. 0.1 kPa (Fig. 4).

### Filmy fern gametophytes are not more stress-tolerant than sporophytes

Contrary to our hypotheses, gametophytes of filmy ferns are in general not more tolerant of the abiotic environment (light levels and water potential) than sporophytes. This differs from other fern groups with desiccation tolerant gametophytes but desiccation avoiding sporophytes (Watkins et al. 2007a). In the filmy ferns we studied, DT levels of both sporophytes and gametophytes largely reflected habitat; in general, epiphytic species tended to have higher DT than terrestrial species, regardless of life stage (Figs 3, S8). Interestingly, this echoes much earlier field observations (not experimental data) by Holloway (1930) on filmy fern gametophytes in New Zealand: he reported apparent DT in gametophytes of *Hymenophyllum rarum* R. Br. and *H. villosum* Colenso, both high elevation epiphytes with desiccation tolerant sporophytes, and lack of DT in terrestrial filamentous gametophytes of *Polyphlebium colensoi* (Hook. f.) Ebihara & K. Iwats., *Abrodictyum strictum* (Menzies ex Hook. & Grev.) Ebihara & K. Iwats., and *A. elongatum* (A. Cunn.) Ebihara & K. Iwats. occupying moist, protected sites on the forest floor. Watkins et al. (2007a) also found that DT of fern gametophytes tended to match habitat, with terrestrial species having lower DT than epiphytes. However, we found that gametophytes had lower DT than sporophytes regardless of growth habit (Figs 3, S8). At high elevation sites (i.e., cloud forest), epiphytic ferns often occur in a dense matrix of bryophytes (Fig. 1b). It is possible that this bryophyte cover helps to retain moisture during brief periods of drought and thus protect the epiphytic fern gametophytes that grow amongst them (Harrington and Watts 2021). More detailed studies of the high elevation epiphytic environment and interactions between co-occurring gametophytes of ferns and bryophytes are required to test this hypothesis.

Although we predicted that species with gametophytes that occur far beyond the range of their sporophytes would have higher DT, this was not the case (Fig. 4c). This is perhaps best illustrated by *C. apiifolia* (Fig. 1d, e, h). Sporophytes of this species are usually restricted to moist cloud forest sites from *c*. 600 m to 1200 m, but we observed gametophytes over a much wider range, down to 200 m (Fig. S3). A similar distribution pattern has been reported in congeneric *Callistopteris baldwinii* (D. C. Eaton) Copel. from Hawaii (Dassler and Farrar 1997). Since gametophytes of *C. apiifolia* are common on Moorea and have a unique morphology (Nitta et al. 2017), we were able to collect sufficient material of this species for additional tests of DT at 100% RH for 2 d. However, *C. apiifolia* gametophytes still failed to recover from even this gentle treatment (Fig. S9). This suggests that the widespread gametophytes of *C. apiifolia* do not rely on DT to attain their distribution, but instead must do so via some other mechanism. Although we measured terrestrial microhabitats by positioning sensors a few cm above the ground, these sensors are still outside of the boundary layer. It is likely that the gametophytes of *C. apiifolia* do not actually experience the conditions measured with our ‘terrestrial’ dataloggers, but instead grow within a more consistently moist boundary layer. Gametophytes of *C. apiifolia* produce gemmae (Fig. 1e, h), and often form dense clonal mats. In the case that their microhabitats do become too dry, they may suffer loss at the edge of the population, then recover by clonal growth (Farrar et al. 2008). Furthermore, the complex surface of clonal mats has been shown to increase the boundary layer and decrease evaporation rates in mosses (Rice et al. 2001), and may function similarly in *C. apiifolia*. Other filmy fern species also produce gemmae, but they are much more frequent in *C. apiifolia*, and the large size of the clonal mats formed by *C. apiifolia* indicate that it may have a high growth rate (J. Nitta, pers. obs.). It is possible that there is a tradeoff between growth rate and DT, such that species capable of rapid asexual growth like *C. apiifolia* are less able to tolerate desiccation, but slower growing epiphytes such as *Hymenophyllum* spp. can better tolerate it (Oliver et al. 2000). Growth rates of tolerant versus non-tolerant gametophytes, including quantification of gemmae production, should be investigated in these species to test this hypothesis.

### Gametophytes are adapted for lower light levels than sporophytes

We observed a clear difference between life stages in photosynthetic optima, with gametophytes consistently adapted for lower light levels than sporophytes (Fig. 3, Table 2). In the only other study to our knowledge that compared photosynthetic rates between gametophytes and sporophytes of a filmy fern, Johnson et al. (2000) also found that gametophytes of *V. speciosa* reached maximum ETR at lower levels than sporophytes: in gametophytes of this species, photosynthesis became saturated at a PPFD of 4–5 µmol·m^-2^·s^-1^ in the field or 30–50 µmol·m^-2^·s^-1^ in the lab vs. 30–50 µmol·m^-2^ ·s^-1^ for sporophytes (lab measurement only). Although we found similar differences in direction between gametophytes and sporophytes, the measurements of Johnson et al. (2000) are considerably lower than what we observed in the field for both Moorean filmy fern gametophytes (PPFD_95%_ *c*. 40– 60 µmol·m^-2^·s^-1^) and sporophytes (PPFD_95%_ *c*. 130–160 µmol·m^-2^·s^-1^). This makes sense in light of the fact that *V. speciosa* is an extremely deep-shade plant, with gametophytes occupying protected rock crevices that receive less than 1 µmol·m^-2^·s^-1^ PPFD for most of the day (Johnson et al. 2000); many of the gametophytes sampled in our study occur in more exposed conditions. It is surprising, however, that we did not observe any consistent differences between species in different habitats, given that light levels should vary between tree trunks and the forest floor, at least on a *c*. 1 m scale. Although the GLMM showed an interaction effect between life stage and growth habit for PPFD_95%_, the model including only life stage was selected for ETR_max_ (Table 2). It is possible that the actual light levels experienced by gametophytes growing in these habitats may not vary significantly if they occupy protected microsites such as bryophyte mats or crevices in rock or bark.

### Influence of phylogeny on traits

Previous studies of filmy fern ecophysiology also found a link between habitat and DT and/or photosynthetic rates (Parra et al. 2009; Proctor 2003, 2012; Saldaña et al. 2014); however, ours is the first to our knowledge to investigate this pattern while taking phylogeny into account. We detected phylogenetic signal in DT (Table 1), suggesting that phylogeny should be considered when interpreting differences in DT between species. In contrast, the lack of phylogenetic signal in photosynthetic traits, coupled with the clear differences in traits between life stages, suggest that sporophytes and gametophytes are adapted for different light levels regardless of clade or growth habit. In a more densely sampled study of morphological evolution in the trichomanoid clade, Dubuisson et al. (2013) identified several morphological changes including reduced stele and root systems associated with the evolution of epiphytic growth within the HE subclade (comprising the genera *Polyphlebium, Didymoglossum, Crepidomanes*, and *Vandenboschia*). It is possible that both reduced morphology and increased DT are parts of an integrated adaptive strategy for epiphytic growth within this subclade of filmy ferns, and have evolved together repeatedly. However, Dubuisson et al. (2013) were unable to determine if these morphological changes had occurred independently, or if there was a single origin at the base of the HE clade followed by multiple losses. Increased sampling of both morphological and physiological traits of additional trichomanoid taxa is needed to distinguish between these competing scenarios.

### Concluding remarks and future directions

In filmy ferns, both sporophyte and gametophyte life stages are poikilohydric; however, this does not necessarily mean they share similar physiological optima. Although filmy fern sporophytes are often of small stature, elevating a leaf even a few cm above the substrate surface means escape from the boundary layer and exposure to substantially greater evapotranspiration. It is likely that even in this clade, wherein the physiological divide between sporophyte and gametophyte is probably the smallest amongst ferns, the two life stages experience substantially different selective pressures. Fossils indicate that some of the earliest vascular plants may have had sporophytes and gametophytes that were both branched and small (Kenrick and Crane 1997), but it is unclear at what point selective pressures began to diverge to favor large, complex sporophytes and small, simple gametophytes. The relevance of plant size to water relations is supported by a study showing increased DT in smaller size classes of the epiphytic fern *Asplenium auritum* Sw. (Testo and Watkins 2012). Future studies of DT in filmy ferns should include juvenile stages of the sporophyte, which are expected to experience similar microhabitats as gametophytes, to better determine the role of microhabitat vs. life stage (and plant size) on DT. The recent application of transcriptomics to filmy fern ecophysiology has begun to reveal the genetic mechanisms underlying DT in these plants (Ostria-Gallardo et al. 2020a, b). Future studies that apply such methods to gametophytes as well as sporophytes across a variety of growth habits are anticipated to provide even greater insight into the ecology and evolution of filmy ferns as well as the transition from desiccation tolerance to avoidance in vascular plants.

## Supporting information

Supplemental Figures (ESM 1)

Supplemental Table (ESM 2)

## Acknowledgments

Members of the Davis Lab and Jonathan Losos provided helpful discussion and comments on drafts. Saad Amer, Ravahere Taputuarai, Tohei Theophilus, and Suzanne Vinnette assisted with fieldwork and experiments. Weston Testo provided advice on settings for the DT test. Two anonymous reviewers provided feedback that improved the manuscript. Staff at the University of California Berkeley’s Richard B. Gump South Pacific Research Station, Moorea, French Polynesia, in particular Valentine Brotherson, Neil Davies, Hinano Teavai-Murphy, and Frank Murphy, provided logistical support for fieldwork. Funding provided in part by the National Science Foundation (Doctoral Dissertation Improvement Grant DEB-1311169 to JHN and CCD), Setup Funds from Harvard University to CCD, American Society of Plant Taxonomists (Research Grant for Graduate Students to JHN), Garden Club of America (Award in Tropical Botany to JHN), Harvard University Herbaria (Fernald Fieldwork Fellowship to JHN), Society of Systematic Biologists (Graduate Student Research Award to JHN), and Systematics Association (Systematics Research Fund to JHN).

