## Supplemental Figures (ESM 1) for "Ecophysiological differentiation between life stages in filmy ferns (Hymenophyllaceae)"

**Electronic supplementary materials: Online Resource 1**

**Content:** Figs. S1–S9

**Fig. S1**. Maximum likelihood ultrametric phylogenetic tree including all filmy fern species from Moorea, French Polynesia.

**Fig. S2**. Temperature, relative humidity (RH), and duration of desiccation treatment of filmy ferns from Moorea, French Polynesia.

**Fig. S3**. Elevational range (m) of sporophytes and gametophytes of filmy ferns on Moorea, French Polynesia.

**Fig. S4**. Recovery of controls (samples kept on moist tissues) during desiccation tolerance test of filmy fern sporophytes from Moorea, French Polynesia.

**Fig. S5**. Relative water content (RWC) of sporophytes after desiccation by duration of desiccation treatment (2 or 15 days).

**Fig. S6**. Relative water content (RWC) of sporophytes during desiccation tolerance test.

**Fig. S7**. Rapid light response curves of filmy ferns from Moorea, French Polynesia.

**Fig. S8**. Box plots comparing physiological parameters between sporophytes (red) and gametophytes (blue) across growth habits in filmy ferns from Moorea, French Polynesia.

**Fig. S9**. Recovery of *Callistopteris apiifolia* gametophytes from Moorea, French Polynesia from desiccation treatment at 100 % RH (water used in place of desiccation salt).

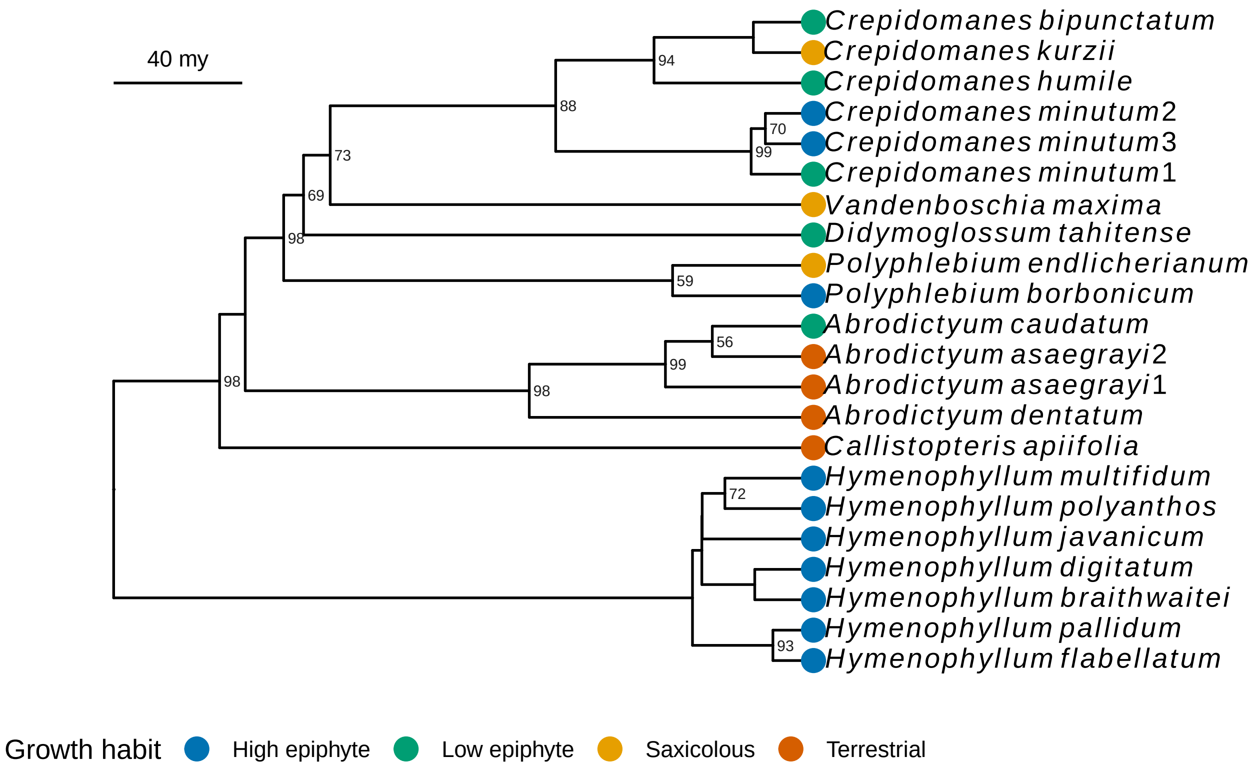


**Fig. S1**. Maximum likelihood ultrametric phylogenetic tree including all filmy fern species from Moorea, French Polynesia. Tree obtained by extracting Hymenophyllaceae from the tree of Nitta et al. (2017) with the 'extract.clade' function of the 'ape' package (Paradis et al. 2004). Bootstrap support values > 50 % shown at nodes. Growth habit indicated by colored dots. *Crepidomanes bipunctatum* and *C. humile* were observed growing as both epiphytes and saxicoles, but are shown as epiphytes to distinguish them from exclusively saxicolous species.

**a**

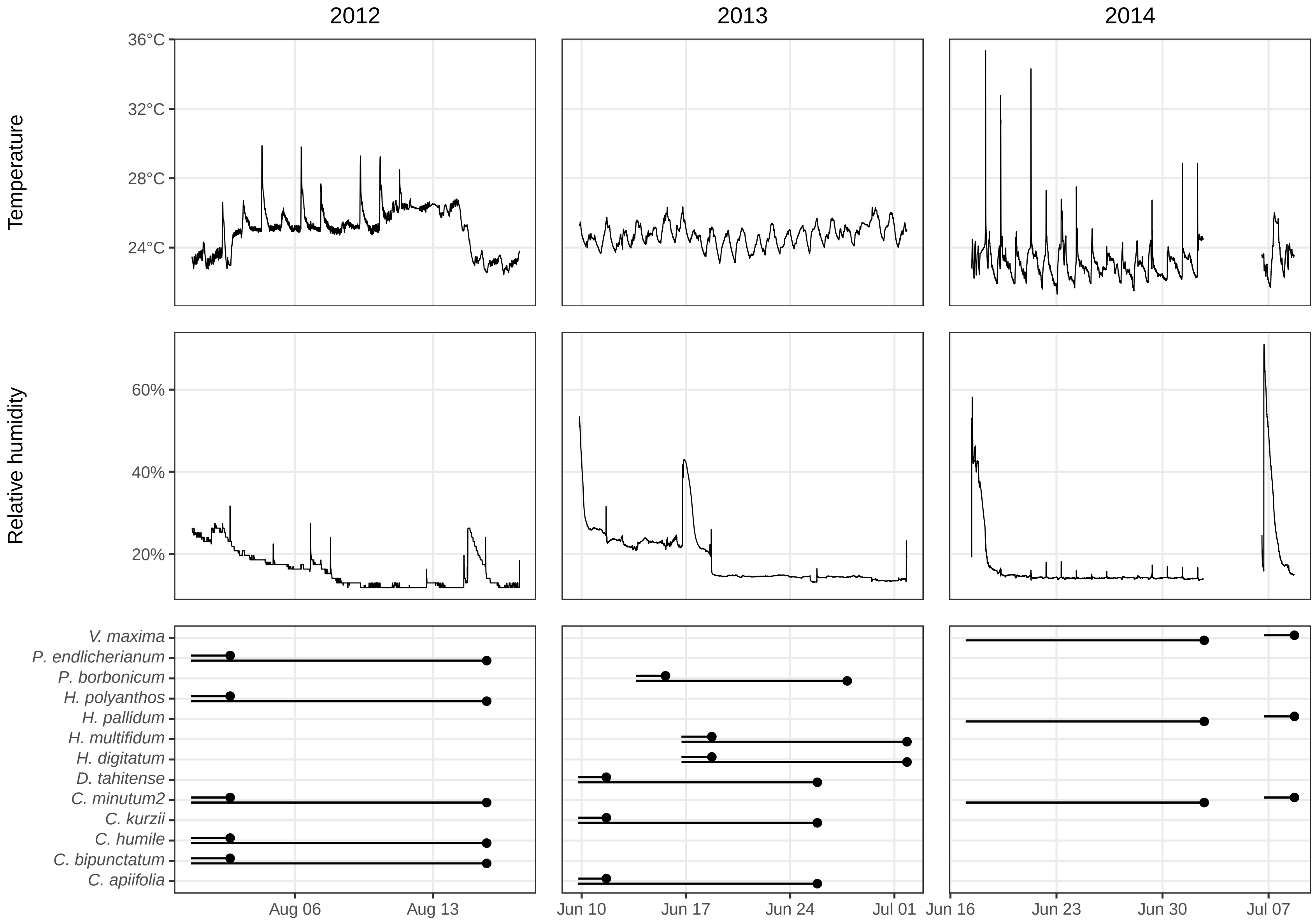
2012
2013
2014
Temperature
36°C
32°C
28°C
24°C
Relative humidity
60%
40%
20%
V. maxima
P. endlicherianum
P. borbonicum
H. polyanthos
H. pallidum
H. multifidum
H. digitatum
D. tahitense
C. minutum2
C. kurzii
C. humile
C. bipunctatum
C. apiifolia
Aug 06
Aug 13
Jun 10
Jun 17
Jun 24
Jul 01
Jun 16
Jun 23
Jun 30
Jul 07

**b**

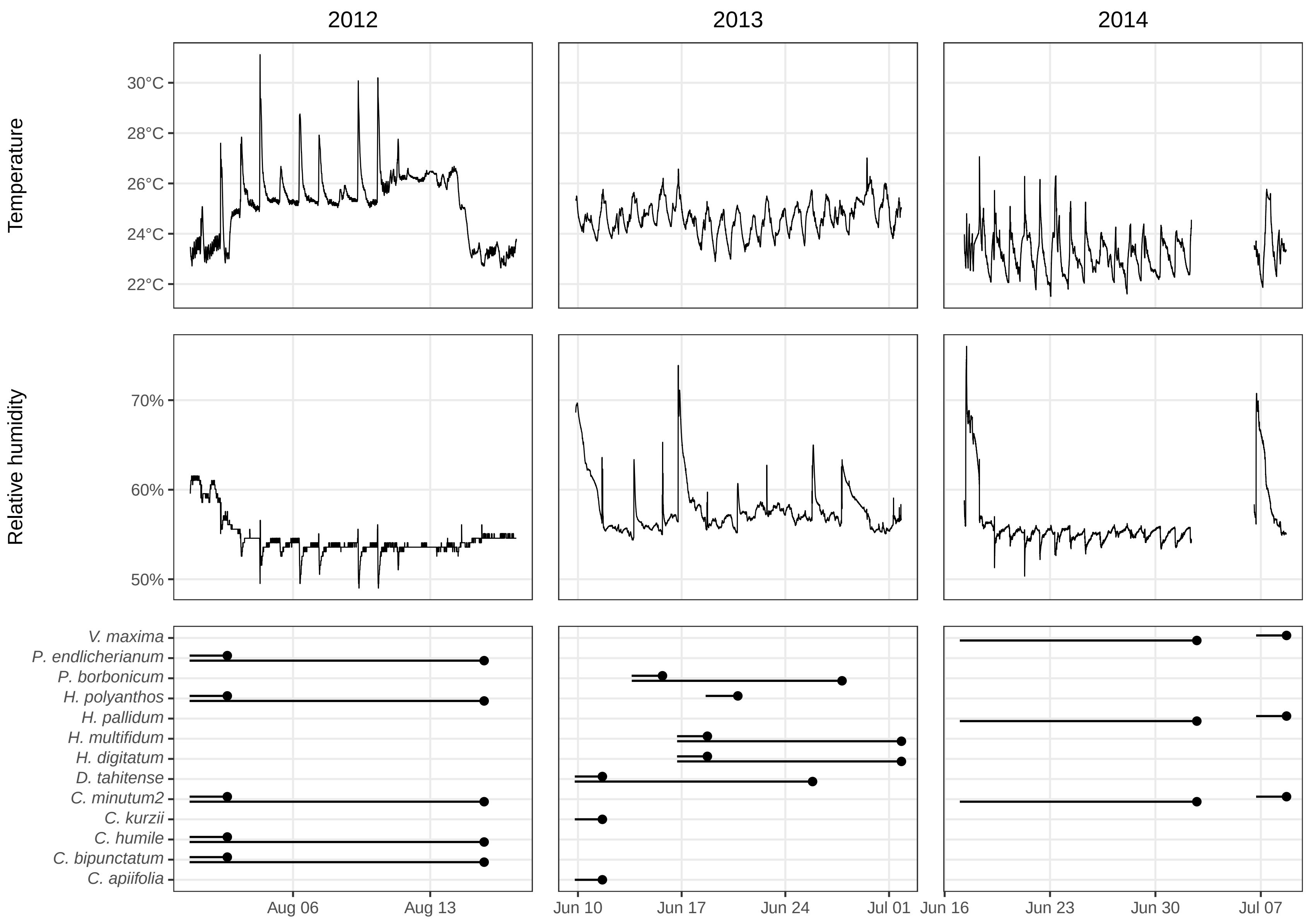
2012
2013
2014
Temperature
30°C
28°C
26°C
24°C
22°C
Relative humidity
70%
60%
50%
V. maxima
P. endlicherianum
P. borbonicum
H. polyanthos
H. pallidum
H. multifidum
H. digitatum
D. tahitense
C. minutum2
C. kurzii
C. humile
C. bipunctatum
C. apiifolia
Aug 06
Aug 13
Jun 10
Jun 17
Jun 24
Jul 01
Jun 16
Jun 23
Jun 30
Jul 07

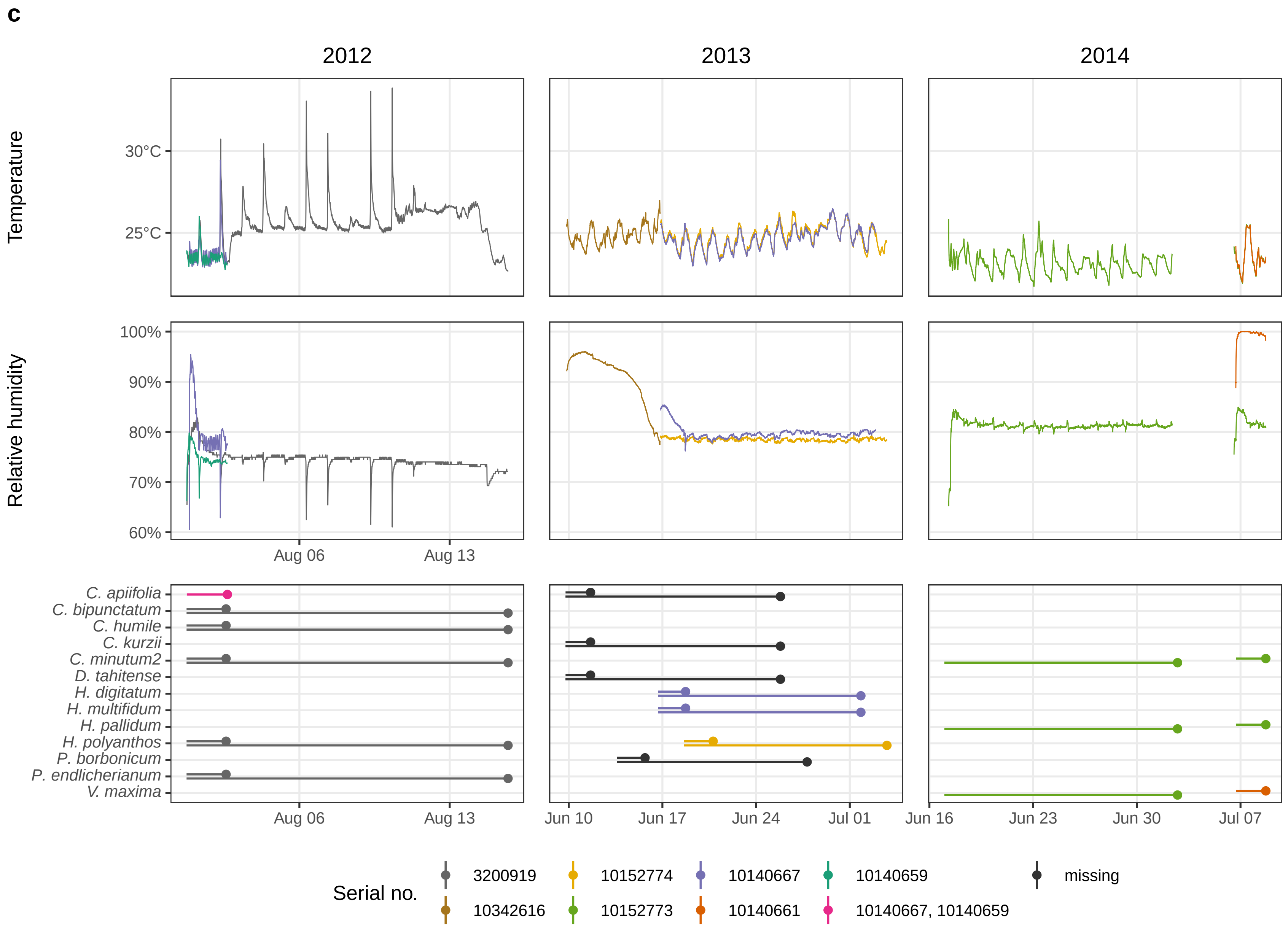
c
2012
2013
2014
Temperature
30°C
25°C
Relative humidity
100%
90%
80%
70%
60%
Aug 06
Aug 13
C. apiifolia
C. bipunctatum
C. humile
C. kurzii
C. minutum2
D. tahitense
H. digitatum
H. multifidum
H. pallidum
H. polyanthos
P. borbonicum
P. endlicherianum
V. maxima
Aug 06
Aug 13
Jun 10
Jun 17
Jun 24
Jul 01
Jun 16
Jun 23
Jun 30
Jul 07
Serial no.
3200919
10152774
10140667
10140659
missing
10342616
10152773
10140661
10140667, 10140659

**d**

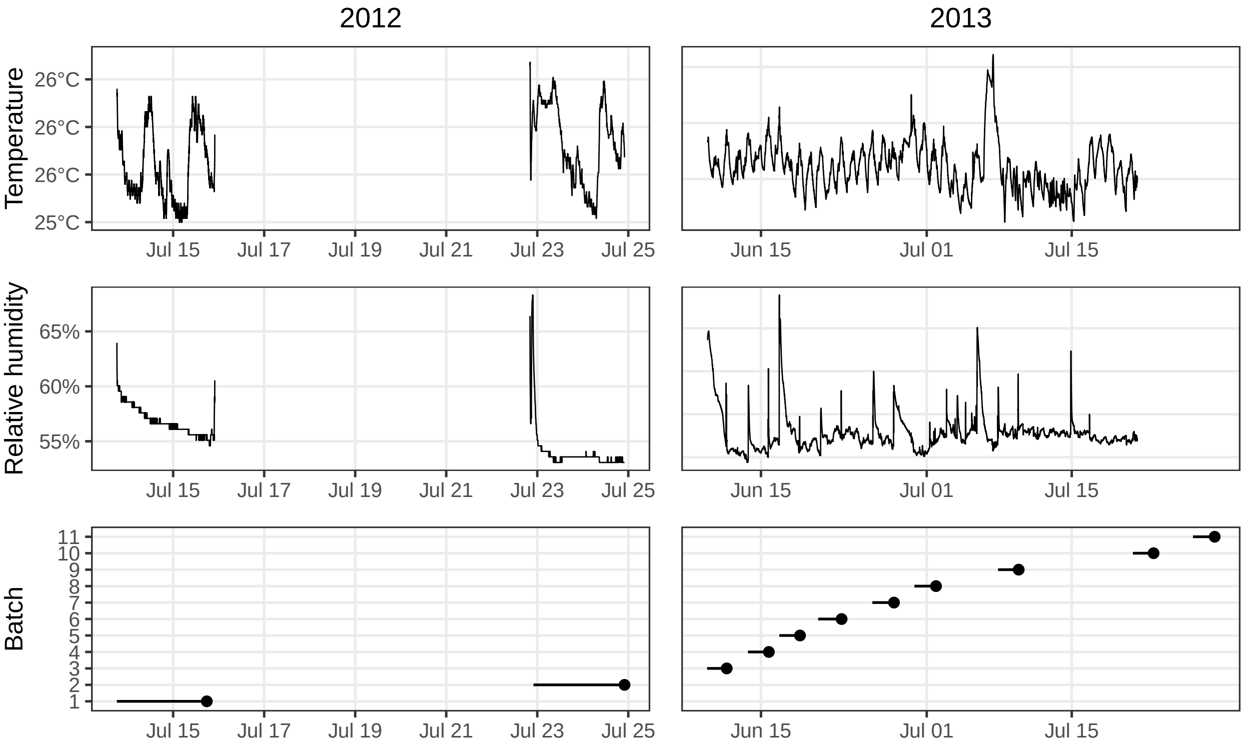
2012
2013
Temperature
26°C
26°C
26°C
25°C
Jul 15
Jul 17
Jul 19
Jul 21
Jul 23
Jul 25
Jun 15
Jul 01
Jul 15
Relative humidity
65%
60%
55%
Batch
11
10
9
8
7
6
5
4
3
2
1

**Fig. S2**. Temperature, relative humidity (RH), and duration of desiccation treatment of filmy ferns from Moorea, French Polynesia. **a**–**c**, sporophytes; **d**, gametophytes. Relative humidity was controlled by placing drying salts inside desiccation chambers: **a**, LiCl (-282 MPa, 18 % RH); **b**, **d**, $Mg(NO_3)_2$ (-86 MPa, 58 % RH); **c**, NaCl (-38 MPa, 80 % RH). Temperature and RH measured with Track-It RH/Temp dataloggers logging every 10 min (2012 species other than *Callistopteris apiifolia*) or Hobo ProV2 dataloggers logging every 5 min (2012 *C. apiifolia*, 2013, 2014). For NaCl (**c**), multiple chambers were set up and tracked with different data loggers; serial number of data logger indicated by color; "missing" indicates correspondence between samples and data loggers (chambers) unknown; *C. apiifolia* samples divided into two chambers. For bottom-most panels (duration of desiccation treatment), line indicates duration of treatment and point indicates removal of sample from chamber. Gametophytes were measured in batches; membership of individual gametophytes to each batch can be found in Table S1 (Online Resource 2).

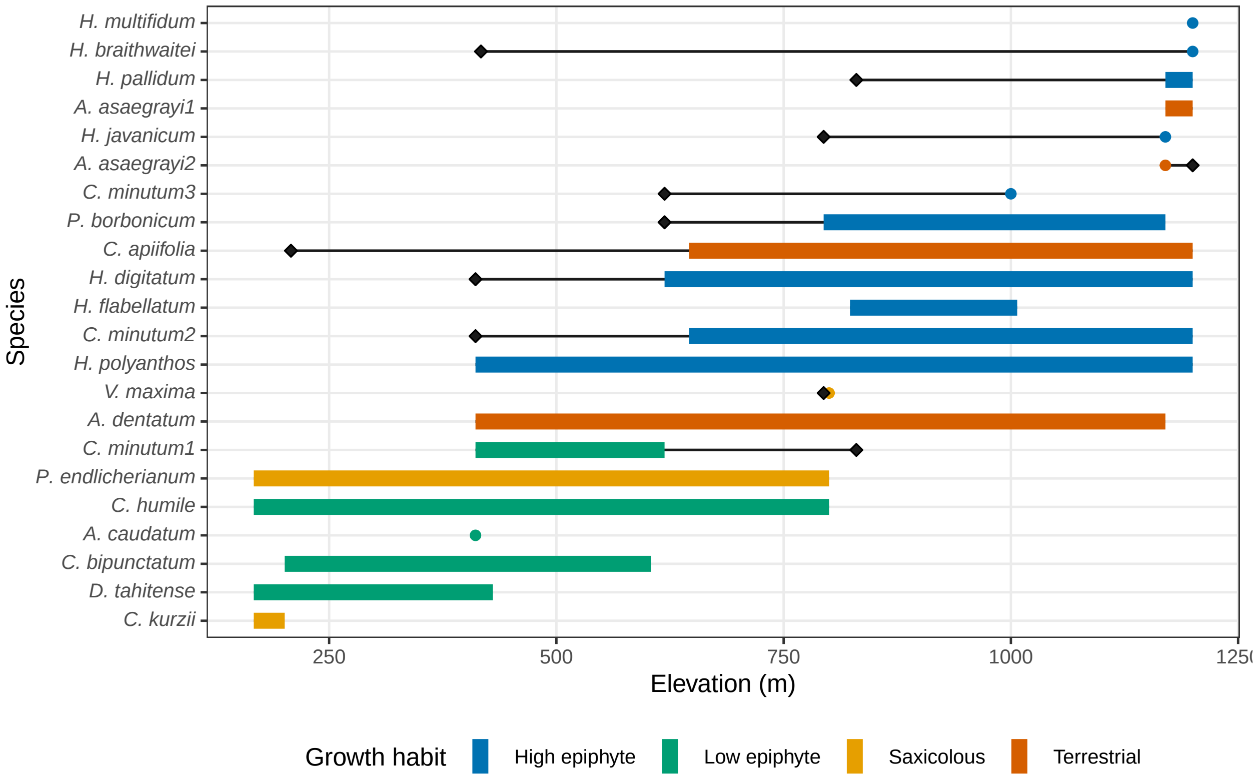


**Fig. S3**. Elevational range (m) of sporophytes and gametophytes of filmy ferns on Moorea, French Polynesia. Elevational range width shown with colored bars (sporophytes) or black lines (gametophytes). Species observed only at a single elevation (not a range) shown with points. Extent of range of gametophytes beyond sporophytes shown with black diamonds. Color indicates growth habit. Presence of gametophytes assumed at sites where sporophytes were observed, even if gametophytes of that species were not collected from that site.

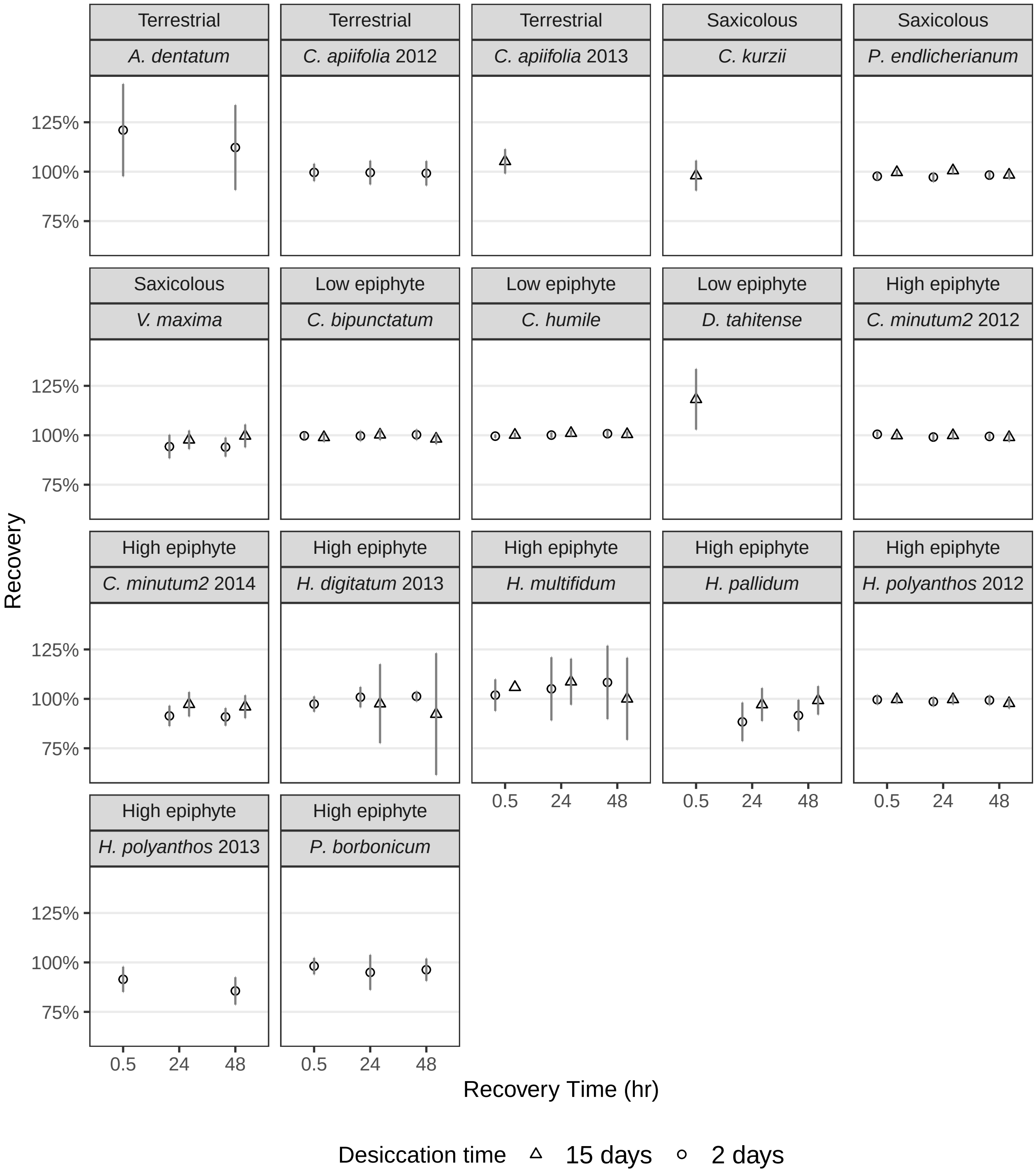


**Fig. S4**. Recovery of controls (samples kept on moist tissues) during desiccation tolerance test of filmy fern sporophytes from Moorea, French Polynesia. Recovery was measured by comparing maximum photochemical yield of photosystem II before desiccation treatment with values at 30 min, 24 hr, and 48 hr following desiccation treatment (see Methods). Recovery was not measured for all combinations of species and desiccation treatments; missing points indicate data not collected (not recovery of 0%). All error bars shown are standard deviation unless otherwise mentioned.

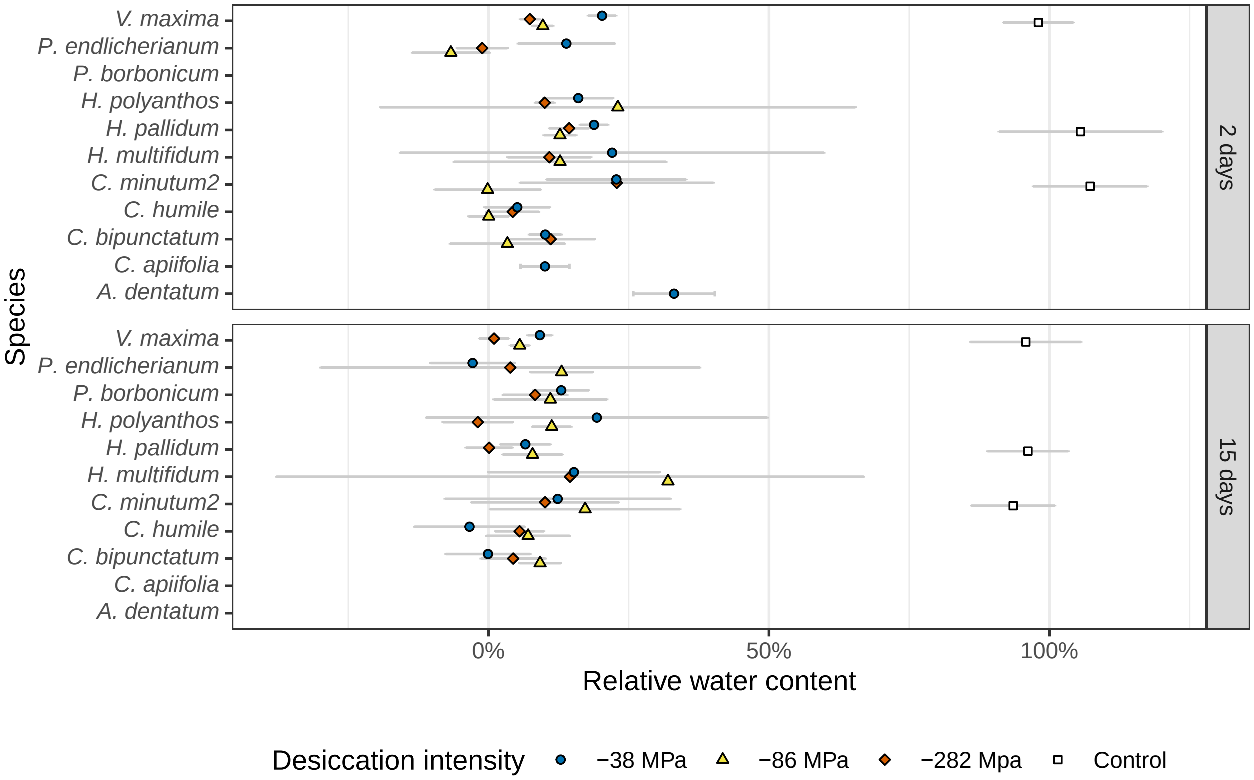


**Fig. S5**. Relative water content (RWC) of sporophytes after desiccation by duration of desiccation treatment (2 or 15 days). RWC was not measured for all combinations of species and desiccation treatments; missing points indicate data not collected (not RWC of zero).

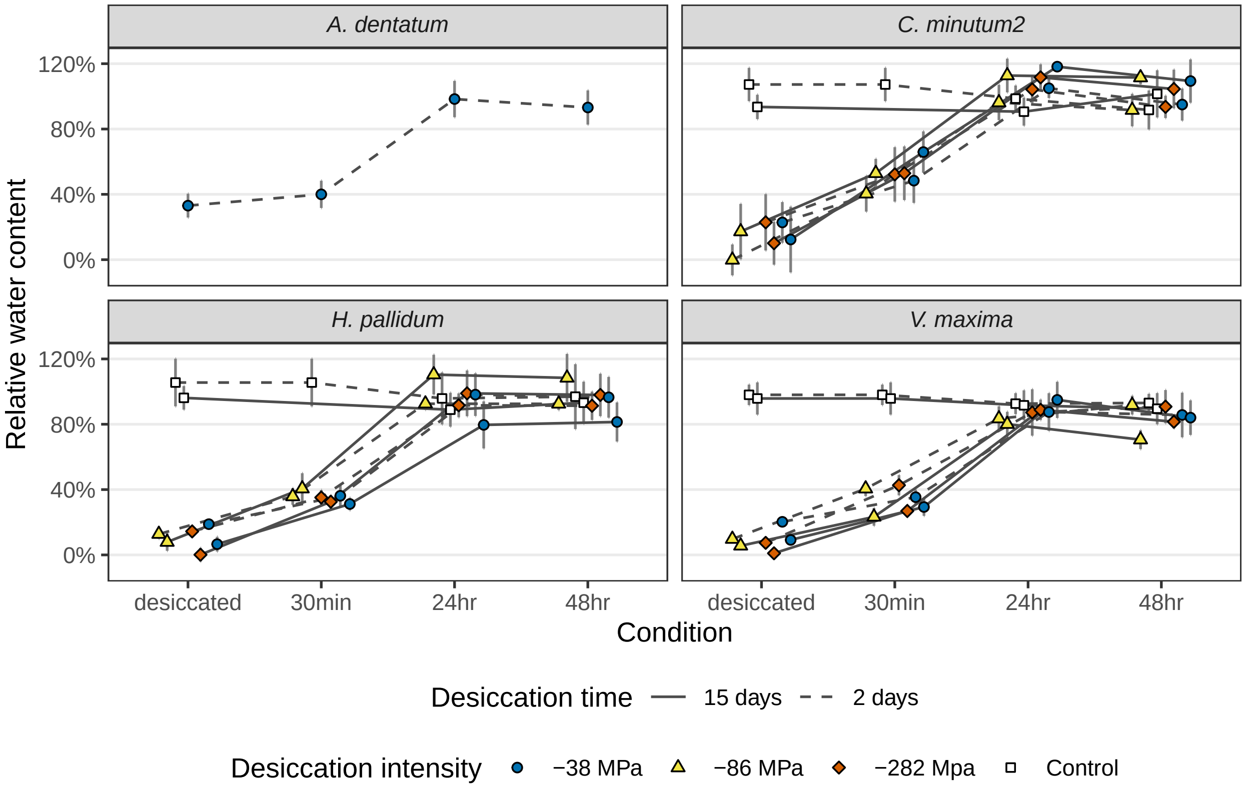


**Fig. S6**. Relative water content (RWC) of sporophytes during desiccation tolerance (DT) test. RWC was not measured for all combinations of species and desiccation treatments; missing points indicate data not collected (not RWC of zero). x-axis indicates condition of sample during DT test; times refer to amount of elapsed time since re-wetting.

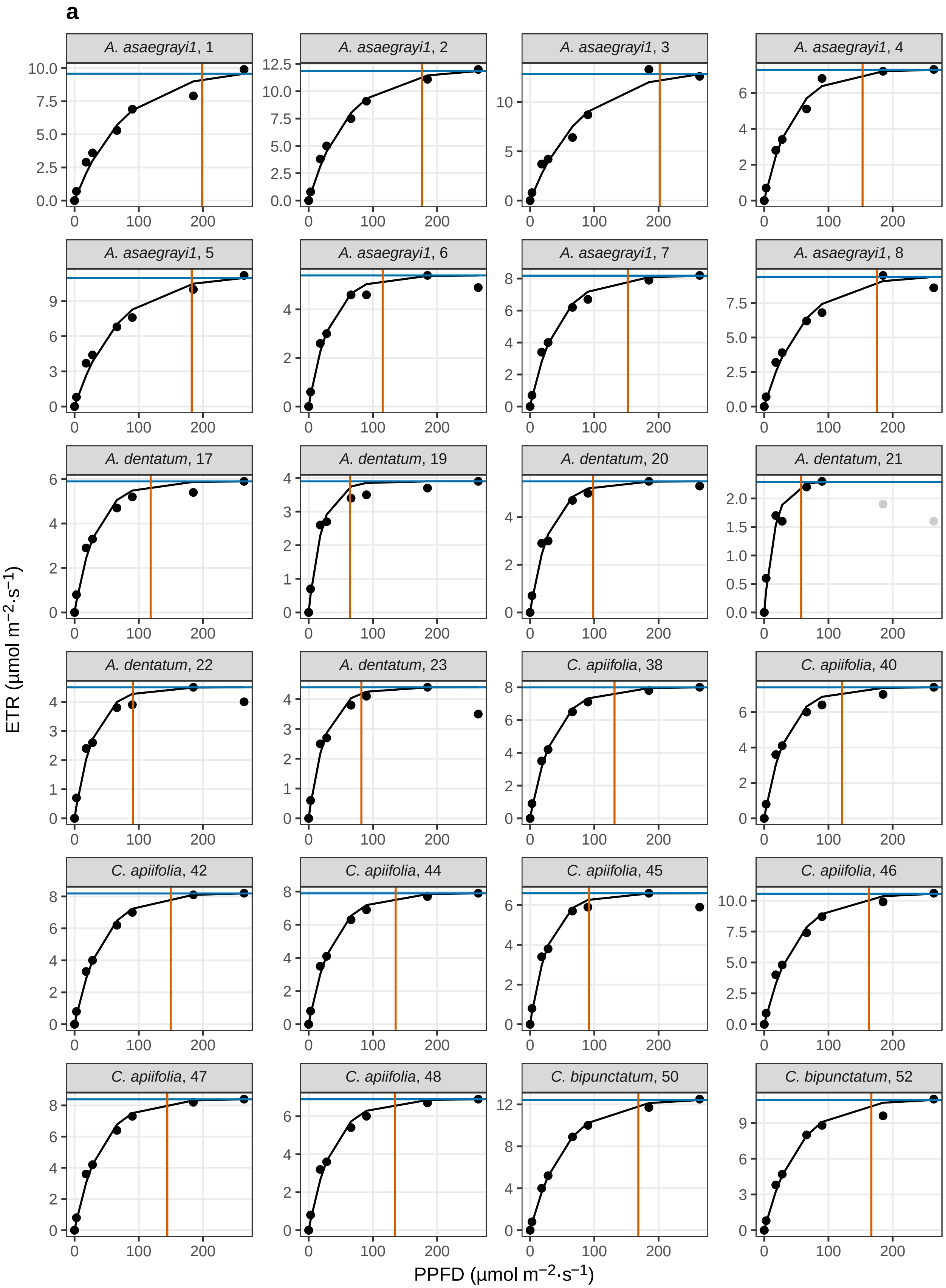

a
A. asaegrayi1, 1
A. asaegrayi1, 2
A. asaegrayi1, 3
A. asaegrayi1, 4
A. asaegrayi1, 5
A. asaegrayi1, 6
A. asaegrayi1, 7
A. asaegrayi1, 8
A. dentatum, 17
A. dentatum, 19
A. dentatum, 20
A. dentatum, 21
A. dentatum, 22
A. dentatum, 23
C. apiifolia, 38
C. apiifolia, 40
C. apiifolia, 42
C. apiifolia, 44
C. apiifolia, 45
C. apiifolia, 46
C. apiifolia, 47
C. apiifolia, 48
C. bipunctatum, 50
C. bipunctatum, 52
ETR (µmol m−2·s−1)
PPFD (µmol m−2·s−1)

**a** (cont.)

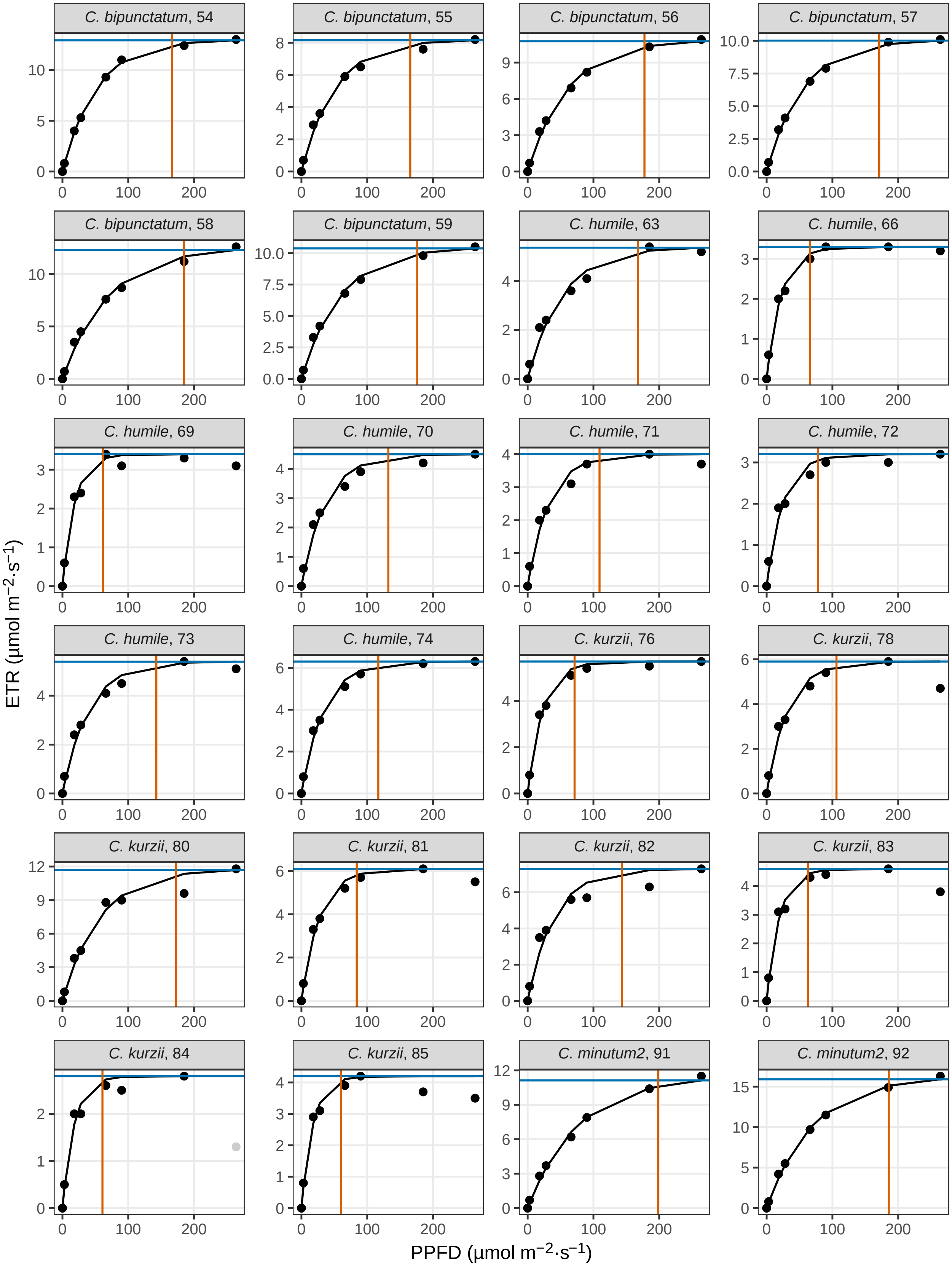
C. bipunctatum, 54
C. bipunctatum, 55
C. bipunctatum, 56
C. bipunctatum, 57
C. bipunctatum, 58
C. bipunctatum, 59
C. humile, 63
C. humile, 66
C. humile, 69
C. humile, 70
C. humile, 71
C. humile, 72
C. humile, 73
C. humile, 74
C. kurzii, 76
C. kurzii, 78
C. kurzii, 80
C. kurzii, 81
C. kurzii, 82
C. kurzii, 83
C. kurzii, 84
C. kurzii, 85
C. minutum2, 91
C. minutum2, 92
ETR (μmol m−2·s−1)
PPFD (μmol m−2·s−1)

**a** (cont.)

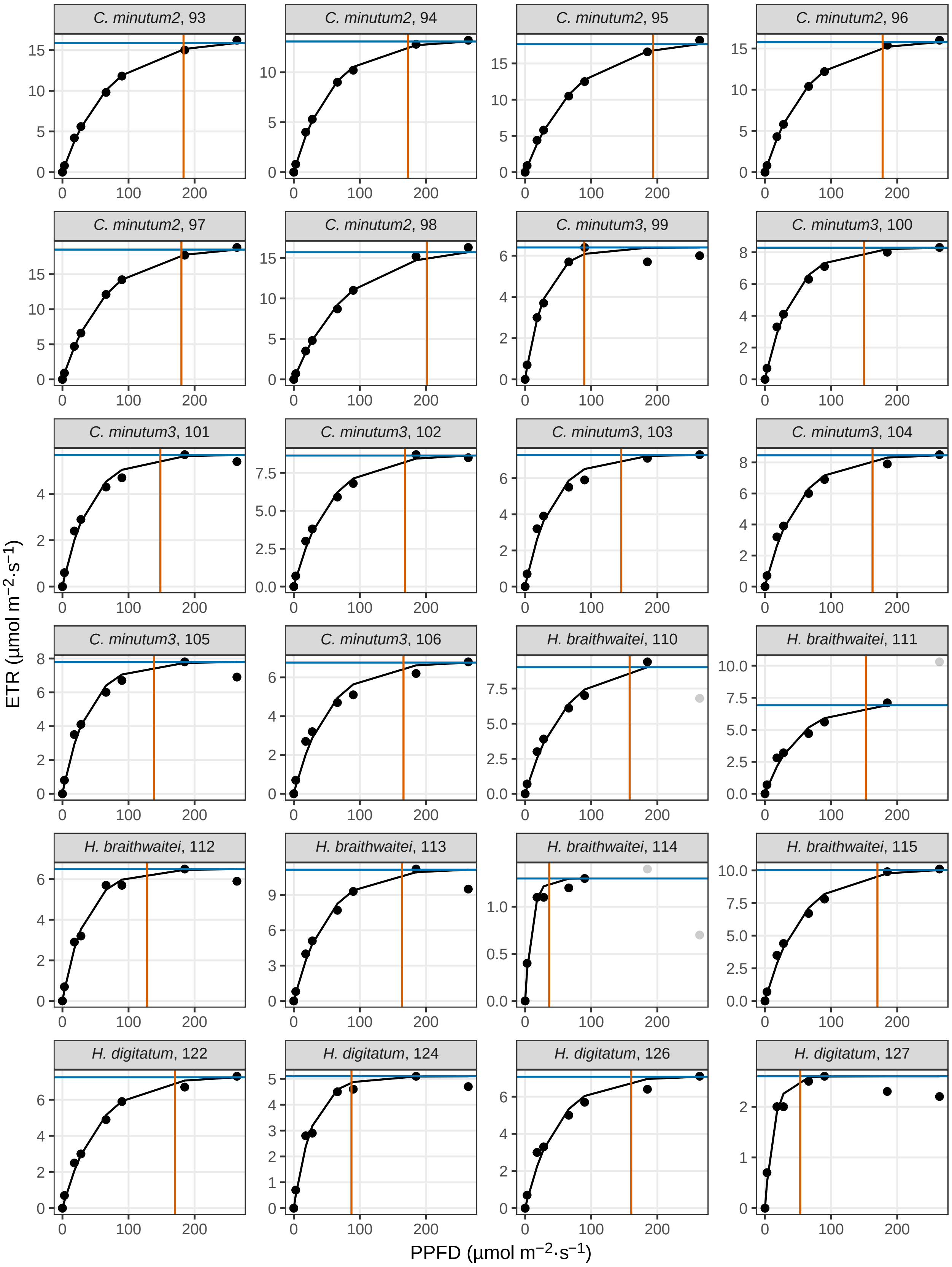
C. minutum2, 93
C. minutum2, 94
C. minutum2, 95
C. minutum2, 96
C. minutum2, 97
C. minutum2, 98
C. minutum3, 99
C. minutum3, 100
C. minutum3, 101
C. minutum3, 102
C. minutum3, 103
C. minutum3, 104
C. minutum3, 105
C. minutum3, 106
H. braithwaitei, 110
H. braithwaitei, 111
H. braithwaitei, 112
H. braithwaitei, 113
H. braithwaitei, 114
H. braithwaitei, 115
H. digitatum, 122
H. digitatum, 124
H. digitatum, 126
H. digitatum, 127
ETR (μmol m−2·s−1)
PPFD (μmol m−2·s−1)

**a** (cont.)

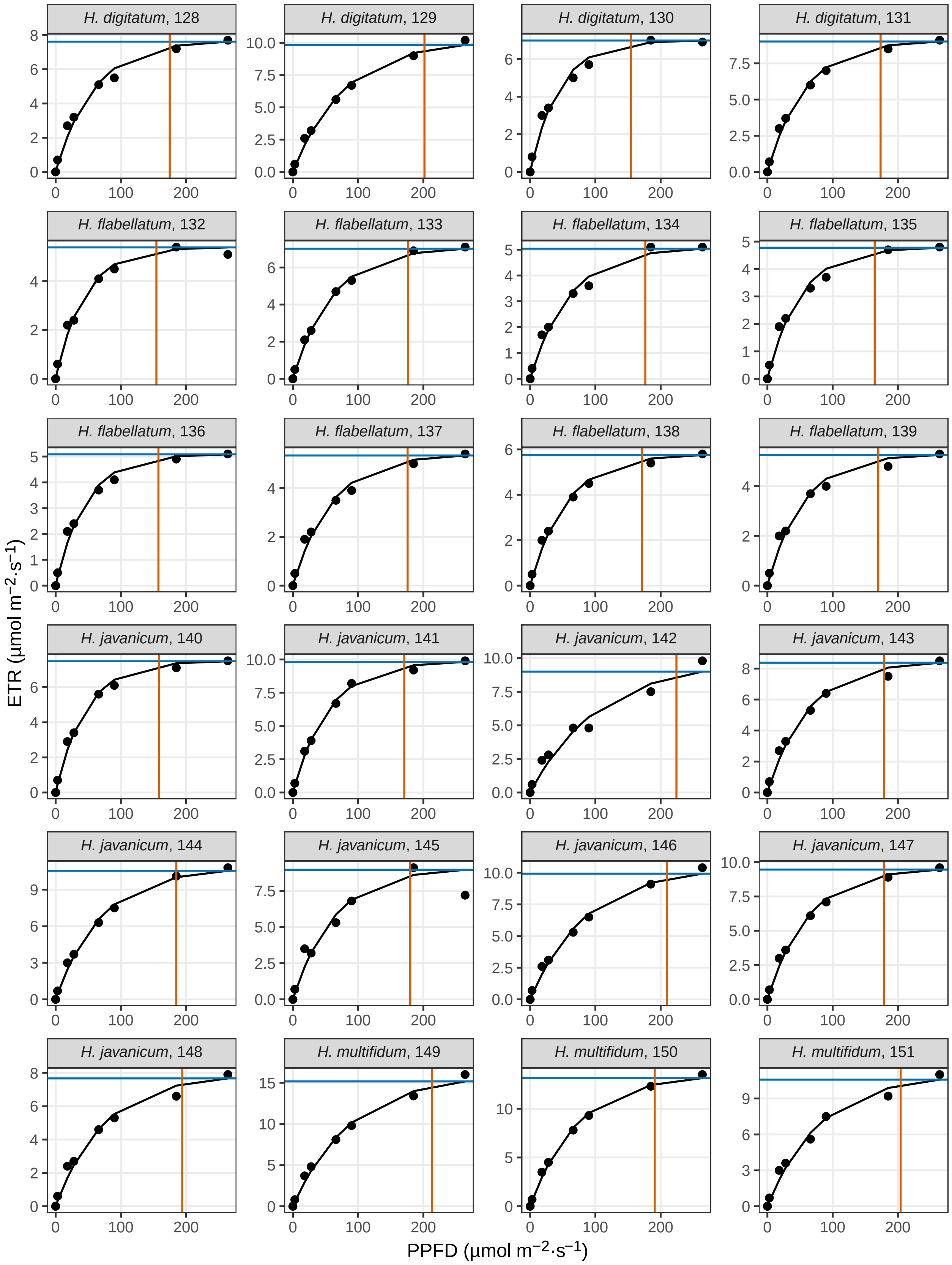
H. digitatum, 128
H. digitatum, 129
H. digitatum, 130
H. digitatum, 131
H. flabellatum, 132
H. flabellatum, 133
H. flabellatum, 134
H. flabellatum, 135
H. flabellatum, 136
H. flabellatum, 137
H. flabellatum, 138
H. flabellatum, 139
H. javanicum, 140
H. javanicum, 141
H. javanicum, 142
H. javanicum, 143
H. javanicum, 144
H. javanicum, 145
H. javanicum, 146
H. javanicum, 147
H. javanicum, 148
H. multifidum, 149
H. multifidum, 150
H. multifidum, 151
ETR (µmol m−2·s−1)
PPFD (µmol m−2·s−1)

**a** (cont.)

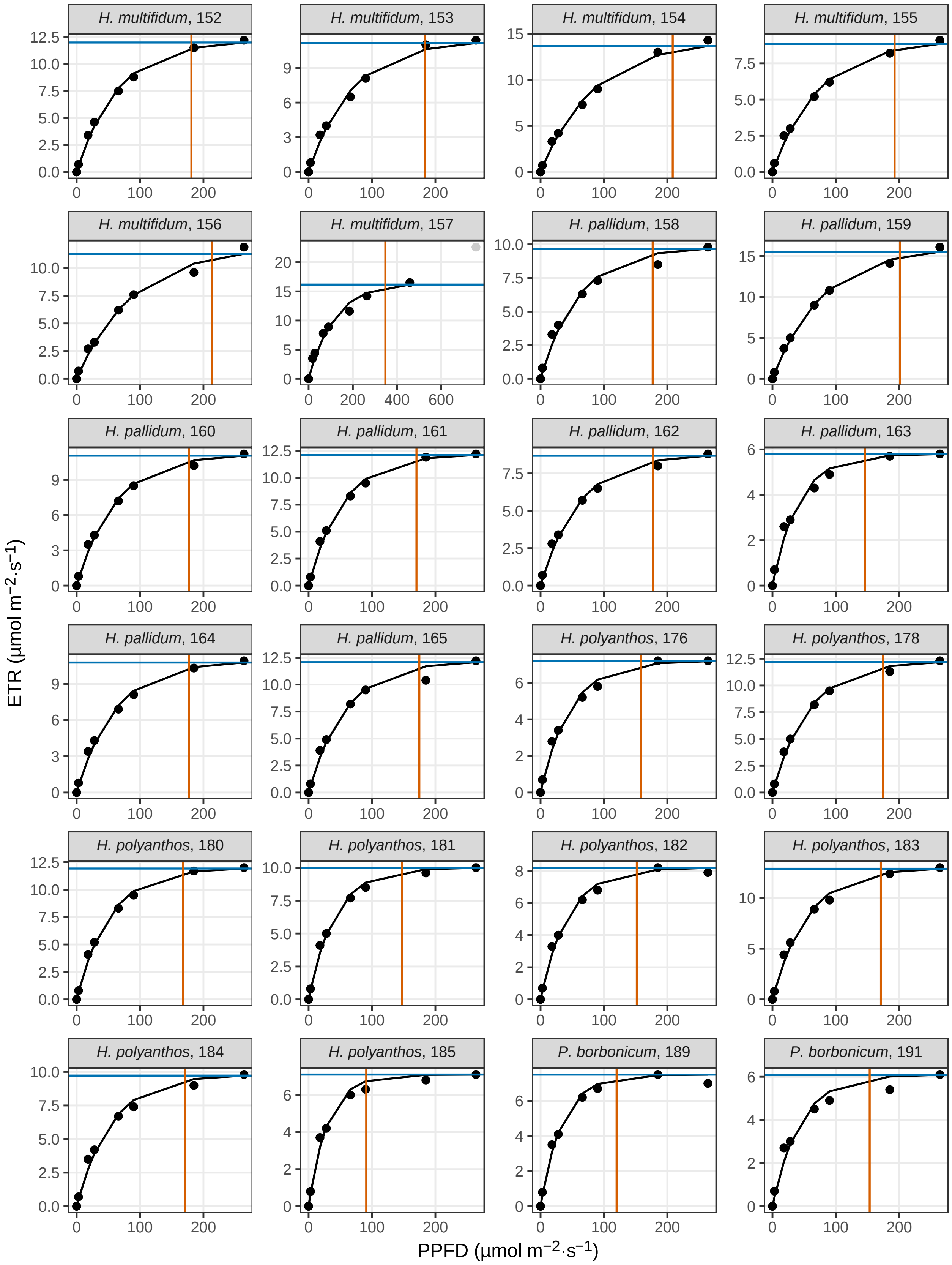
H. multifidum, 152
H. multifidum, 153
H. multifidum, 154
H. multifidum, 155
H. multifidum, 156
H. multifidum, 157
H. pallidum, 158
H. pallidum, 159
H. pallidum, 160
H. pallidum, 161
H. pallidum, 162
H. pallidum, 163
H. pallidum, 164
H. pallidum, 165
H. polyanthos, 176
H. polyanthos, 178
H. polyanthos, 180
H. polyanthos, 181
H. polyanthos, 182
H. polyanthos, 183
H. polyanthos, 184
H. polyanthos, 185
P. borbonicum, 189
P. borbonicum, 191
ETR (μmol m−2·s−1)
PPFD (μmol m−2·s−1)

**a** (cont.)

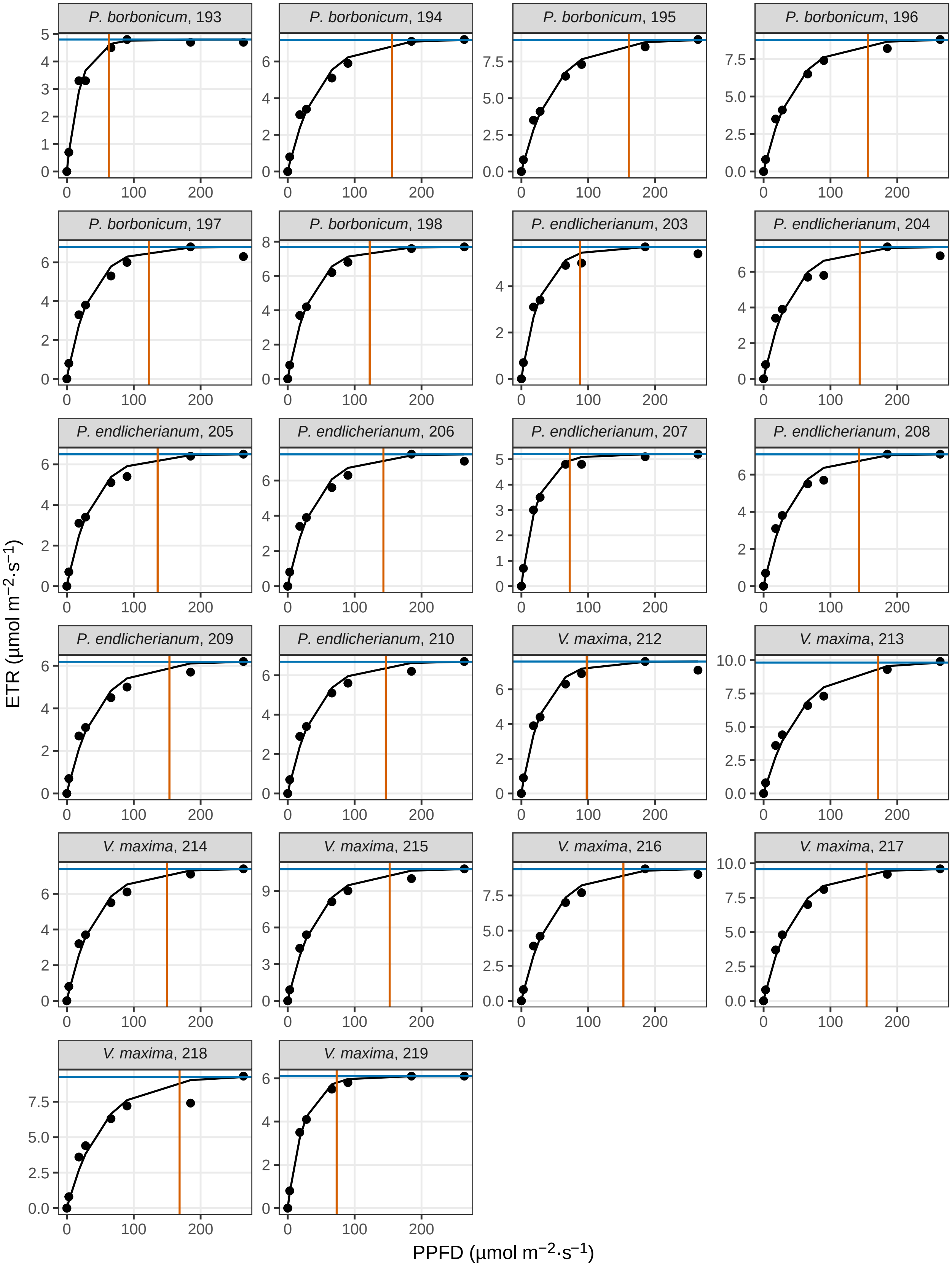

P. borbonicum, 193
P. borbonicum, 194
P. borbonicum, 195
P. borbonicum, 196
P. borbonicum, 197
P. borbonicum, 198
P. endlicherianum, 203
P. endlicherianum, 204
P. endlicherianum, 205
P. endlicherianum, 206
P. endlicherianum, 207
P. endlicherianum, 208
P. endlicherianum, 209
P. endlicherianum, 210
V. maxima, 212
V. maxima, 213
V. maxima, 214
V. maxima, 215
V. maxima, 216
V. maxima, 217
V. maxima, 218
V. maxima, 219
ETR (µmol m−2·s−1)
PPFD (µmol m−2·s−1)

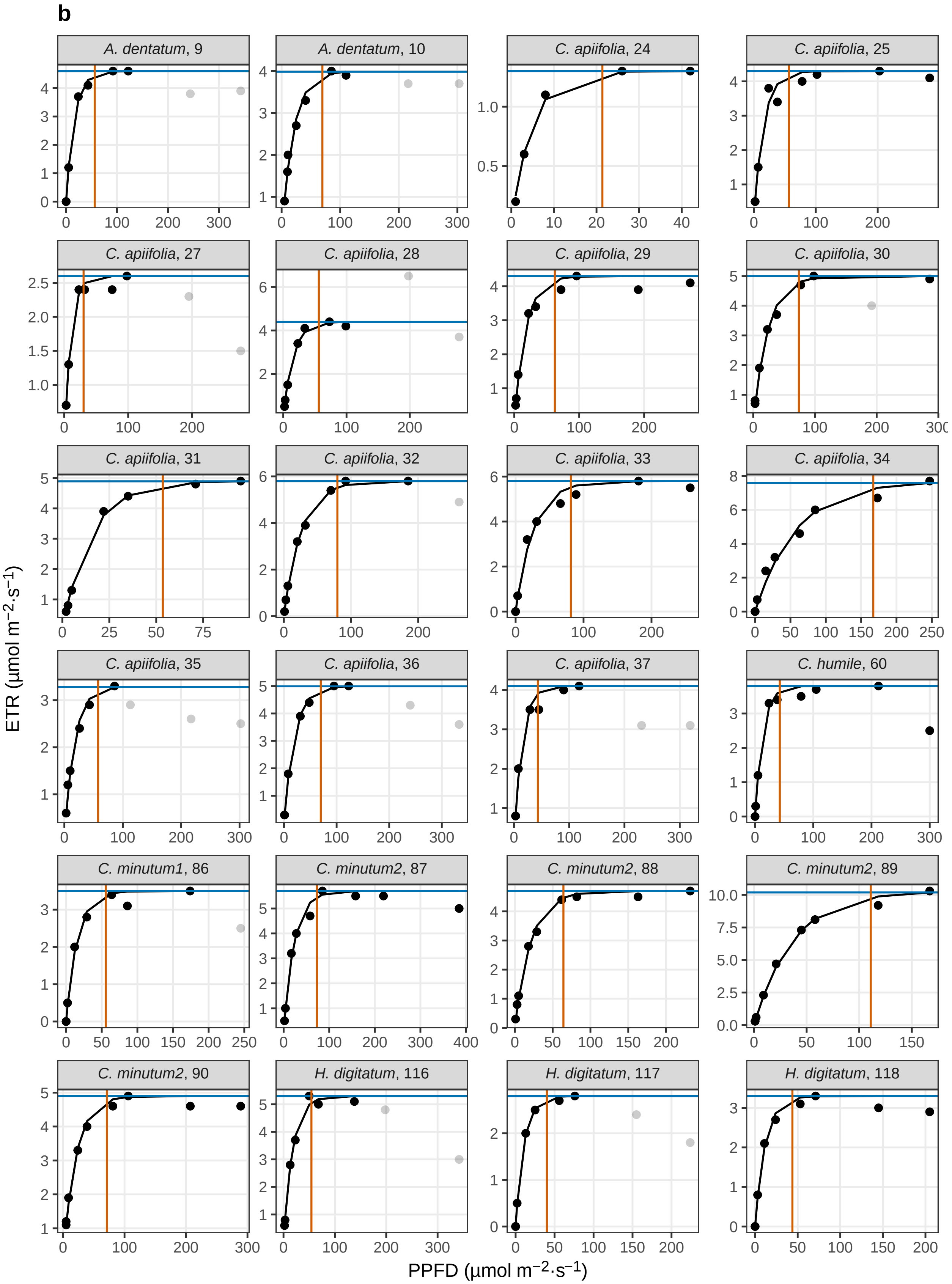
b
A. dentatum, 9
A. dentatum, 10
C. apiifolia, 24
C. apiifolia, 25
C. apiifolia, 27
C. apiifolia, 28
C. apiifolia, 29
C. apiifolia, 30
C. apiifolia, 31
C. apiifolia, 32
C. apiifolia, 33
C. apiifolia, 34
C. apiifolia, 35
C. apiifolia, 36
C. apiifolia, 37
C. humile, 60
C. minutum1, 86
C. minutum2, 87
C. minutum2, 88
C. minutum2, 89
C. minutum2, 90
H. digitatum, 116
H. digitatum, 117
H. digitatum, 118
ETR (µmol m⁻²·s⁻¹)
PPFD (µmol m⁻²·s⁻¹)

**b** (cont.)

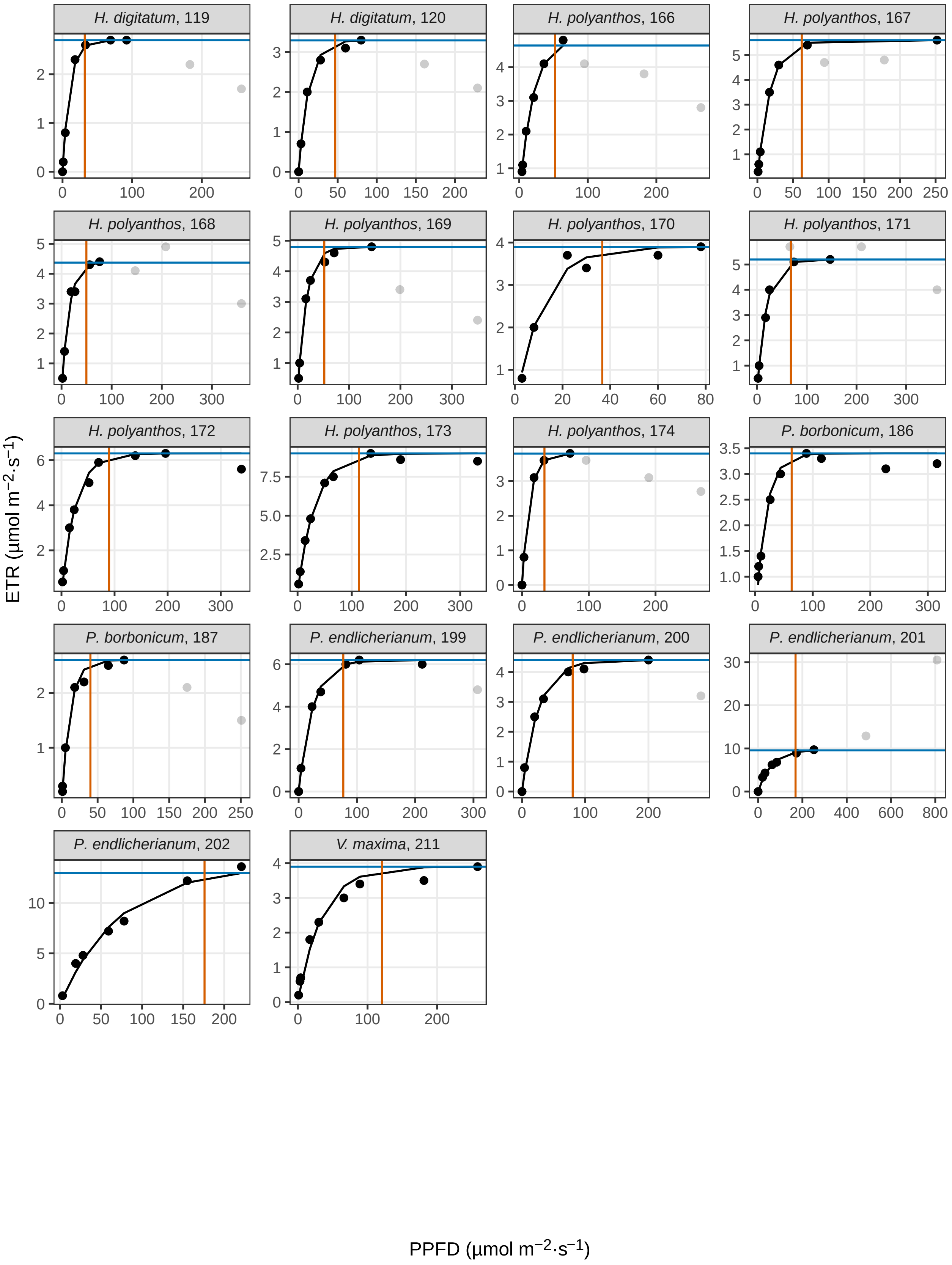

H. digitatum, 119
H. digitatum, 120
H. polyanthos, 166
H. polyanthos, 167
H. polyanthos, 168
H. polyanthos, 169
H. polyanthos, 170
H. polyanthos, 171
H. polyanthos, 172
H. polyanthos, 173
H. polyanthos, 174
P. borbonicum, 186
P. borbonicum, 187
P. endlicherianum, 199
P. endlicherianum, 200
P. endlicherianum, 201
P. endlicherianum, 202
V. maxima, 211
ETR (µmol m−2·s−1)
PPFD (µmol m−2·s−1)

**Fig. S7**. Rapid light response curves of filmy ferns from Moorea, French Polynesia. **a**, sporophytes; **b**, gametophytes. Title of each subplot indicates species followed by light curve ID number assigned to each sample (light curve IDs match those in filmy_light_curves.csv data file). Light curves fit using the equation $y = A(1 - e^{-kx})$, where *y* is relative electron transport rate of photosystem II (ETR), *x* is photosynthetic photon flux density (PPFD), *A* is the asymptote of the curve, and *k* is a slope parameter. Outliers at high PPFD not included in the model indicated in light grey. Blue line indicates maximum modeled relative electron transport rate ($ETR_{max}$). Red line indicates PPFD at 95 % of $ETR_{max}$ ($PPFD_{95\%}$).

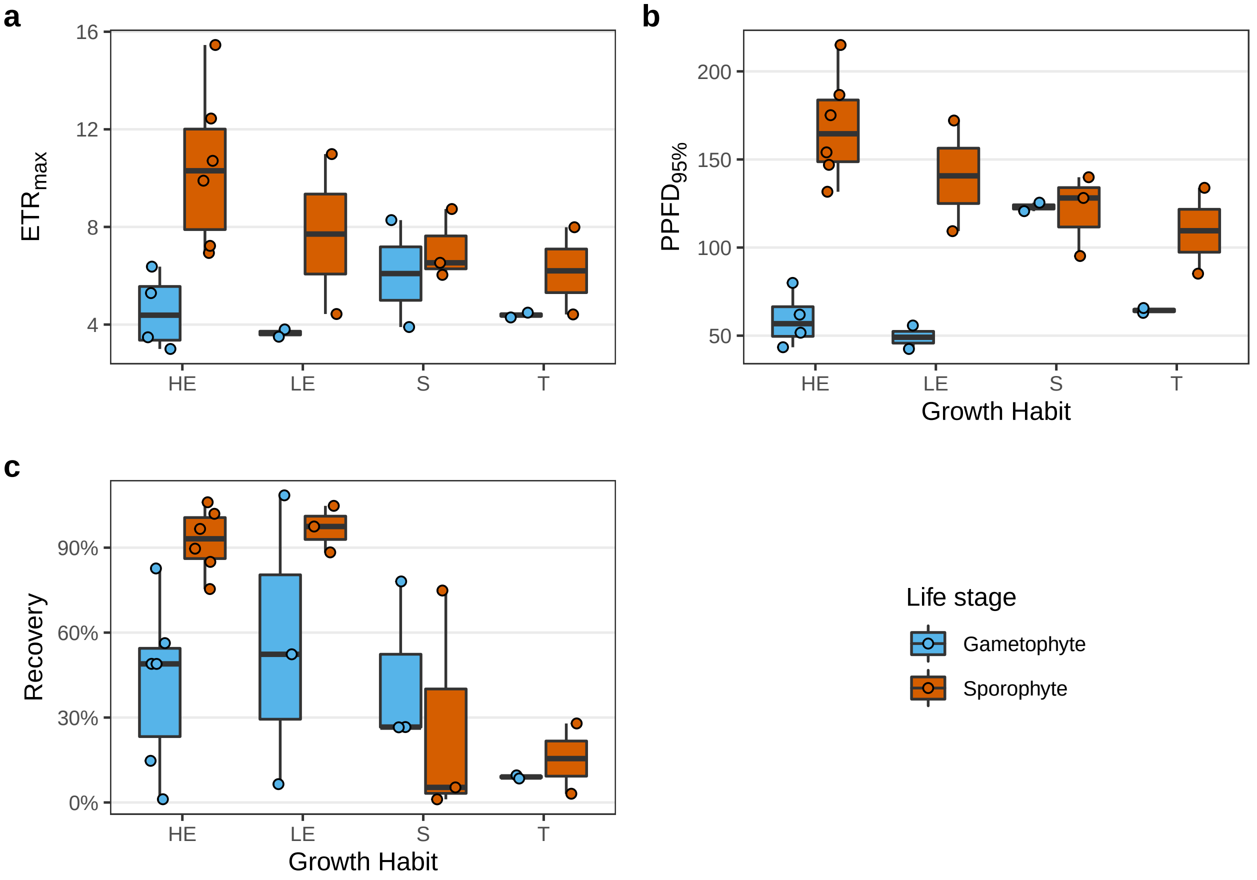


**Fig. S8**. Box plots comparing physiological parameters between sporophytes (red) and gametophytes (blue) across growth habits in filmy ferns from Moorea, French Polynesia. Each point represents one species; dark line is median, lower and upper edges correspond to first and third quartiles, and whiskers extend to the most extreme value no further than 1.5 times the inter-quartile range. **a**, $ETR_{max}$ (μmol electrons $m^{-2} \cdot s^{-1}$); **b** $PPFD_{95\%}$ (μmol photons $m^{-2} \cdot s^{-1}$); **c**, recovery (%) of chlorophyll fluorescence ($F_v/F_m$) following 2 day desiccation at −86 MPa (*Abrodictyum dentatum* was desiccated at −38 MPa instead of −86 MPa). *Crepidomanes bipunctatum* and *C. humile* were observed growing as both epiphytes and saxicoles, but are treated as epiphytes to distinguish them from exclusively saxicolous species. Growth habit abbreviations: HE, high elevation epiphyte; LE, low elevation epiphyte; S, saxicolous; T, terrestrial.

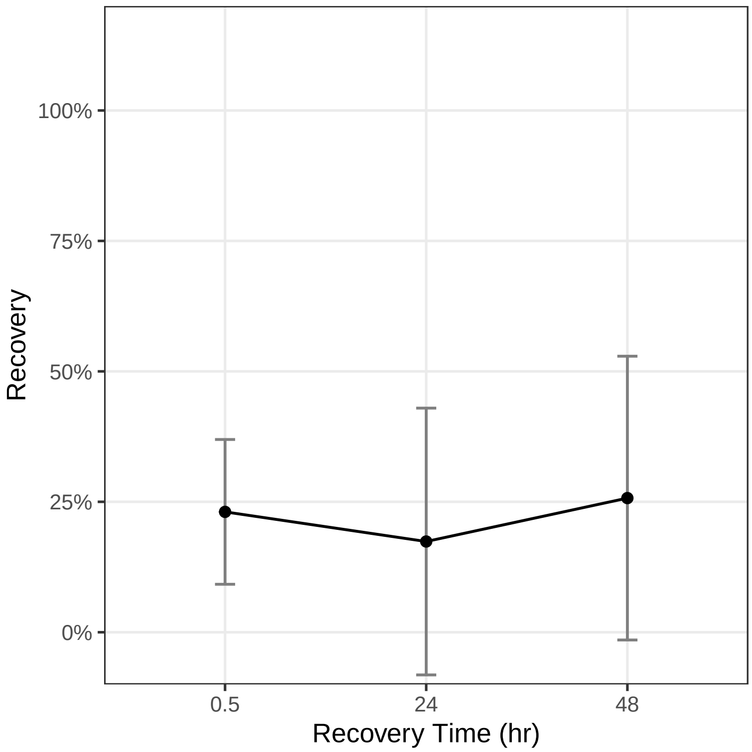


**Fig. S9**. Recovery of *Callistopteris apiifolia* gametophytes from Moorea, French Polynesia from desiccation treatment at 100 % RH (water used in place of desiccation salt). *n* = 7.
